# Assessing the activity of different plant-derived molecules and potential biological nitrification inhibitors on a range of soil ammonia- and nitrite- oxidizing strains

**DOI:** 10.1101/2023.07.12.548655

**Authors:** Maria Kolovou, Dimitra Panagiotou, Lars Süße, Olivier Loiseleur, Simon Williams, Dimitrios G. Karpouzas, Evangelia S. Papadopoulou

## Abstract

Nitrification is associated with significant losses of fertilizer-derived ammonium from agroecosystems. The use of biological nitrification inhibitors (BNIs), in place of synthetic nitrification inhibitors (SNIs), holds a great potential to effectively reduce N losses and conforms with the current move towards ecological-intensified agriculture. Knowledge of the activity of BNIs to soil nitrifiers is limited and is mostly based on bioassays with a single *Nitrosomonas europaea* strain. We determined the *in vitro* activity of multiple plant-derived compounds as BNIs like (i) root-derived compounds (sakuranetin, MHPP, and zeanone); (ii) other phytochemicals (caffeic acid, quinic acid, chlorogenic acid and shikimic acid), and (iii) analogues of statins (simvastatin), triazoles (1-butyl-4-propyl-triazole, 1,4-dibutyltriazole) and zeanone (2-methoxy-1,4-naphthoquinone), on ecophysiologically and phylogenetically distinct soil-derived ammonia-oxidizing bacteria (AOB) (*Nitrosospira multiformis* and *N. europaea*), ammonia-oxidizing archaea (AOA) (*Candidatus* Nitrosotalea sinensis and *Candidatus* Nitrosocosmicus franklandianus), and a nitrite-oxidizing bacterium (NOB) (*Nitrobacter* sp. NHB1). AOA were more sensitive than AOB to BNIs. Sensitivity within AOA group was BNI-dependent, unlike AOB for which *N. multiformis* was consistently more sensitive than *N. europaea*. Interestingly, several compounds were inhibitory to *Nitrobacter* sp. with MHPP and caffeic acid being more potent to the NOB compared to the AOB and AOA strains, an observation with potentially serious implications for soil quality and agricultural production. Overall, zeanone, MHPP and caffeic acid were the most potent ΒNIs towards AOB, zeanone and 2-methoxy-1,4-naphthoquinone were the most effective compounds against AOA, while caffeic acid was the most potent BNI on *Nitrobacter* sp. We provide pioneering evidence for the activity range of multiple BNIs on soil nitrifiers, stress the need for revisiting the biological screening systems currently used for BNI determination and we sought for a more thorough monitoring of the impact of BNI candidates on a range of both target and non-target microorganisms.

## 1. Introduction

Nitrogen (N) is an essential nutrient for plant growth and productivity and increases in its supply through fertilization are actively exploited in agriculture to enhance crop yields (Fowler et al., 2018). Approximately two percent of the world’s energy is consumed for the production of N fertilizers annually (Wendeborn, 2020) with the amount of reactive N (Nr) introduced into the biosphere by these means reaching to 1.5 × 10^8^ t yr^−1^ (Galloway et al., 2008). Critically, due to the low N fertilizer use efficiency (NUE) in agriculture only 30–50% of this amount is assimilated by plants, while the remaining in the forms of ammonia (NH_3_), nitrate (NO_3_^−^), and N oxides (N_x_O), enriches Nr pools and contributes to environmental quality degradation, biodiversity loss, and greenhouse gas (GHG) driven climate change (Cassman et al. 2002; Sutton et al., 2011). N losses from fertilized soils are linked to the microbially mediated process of nitrification, which generates NO_3_^−^, that is mobile in soil and susceptible to groundwater leaching, and it is also a substrate for soil nitrifying and denitrifying microorganisms that could emit nitrous oxide (N_2_O) (Coskun et al. 2017), a GHG with a warming potential 298 times greater than carbon dioxide (Forster et al. 2007).

Currently the use of chemical-based technologies holds a great promise to increase NUE and reduce the adverse environmental impact of Nr cascade. Such chemicals are small molecules (≤ 500 dalton), either synthetic or naturally produced, which can decelerate the key steps of the nitrification process, and particularly the first and often rate-limiting step of ammonia oxidation (Coskun et al. 2017, Qiao et al., 2015; Papadopoulou et al., 2020). In this step, NH_3_ is oxidized to hydroxylamine, by the enzyme ammonia monooxygenase (AMO), and further to nitrite (NO_3_^−^) via nitric oxide (Caranto and Lancaster, 2017), by canonical ammonia-oxidizing bacteria (AOB), ammonia-oxidizing archaea (AOA), and complete ammonia-oxidizing (comammox) bacteria (Daims et al., 2015).

To this end, synthetic nitrification inhibitors (SNIs) like 2-chloro-6-(trichloromethyl) pyridine (nitrapyrin), dicyandiamide (DCD), and 3,4-dimethylpyrazole phosphate (DMPP) are routinely incorporated into fertilizers to stabilize the supply of N available in soil (Subbarao et al., 2006a). The use of SNIs is recommended by the Intergovernmental Panel on Climate Change (IPCC) as a means of mitigating climate change (Gao and Cabrera Serrenho, 2023). However, these compounds have been also reported to have drawbacks such as difficulties in application, highly variable efficacy across soils, high cost, pollution of natural resources, bioaccumulation in plants and animals and potential entry into the food chain (Akiyama et al., 2010; Qiu et al., 2015; Coskun et al. 2017). In response to the constraints of SNIs and to the public demand for novel bio-based solutions in agriculture, biological NIs (BNIs), compounds that are synthesized or secreted by plants and have nitrification inhibition properties, have recently received increased attention as safer and potentially more effective alternatives to SNIs (Wang et al., 2021).

BNI has been thought of as an evolutionary adaptation of certain plants to retain N in the form of NH_4_^+^ in N-limited soils, leading to optimized NUE and a less leaky N cycle (Subbarao et al., 2007). Until now, multiple BNI compounds exudated from the roots of several plant species, mostly grasses, have been identified including sorgoleone, sakuranetin, and methyl 3-(4-hydroxyphenyl) propionate (MHPP) from sorghum (*Sorghum bicolor*) (Subbarao et al., 2013; Zakir et al., 2008), brachialactone from signal grass (*Brachiaria humidicola*) (Subbarao et al., 2009), 1,9-decanediol from rice (*Oryza sativa*) (Sun et al., 2016), 2,7-dimethoxy-1,4-naphthoquinone (zeanone) and 6-methoxy-2(3H)-benzoxazolone (MBOA) from maize (*Zea mays*) (Otaka et al., 2022; 2023). BNIs can be also plant secondary metabolites (e.g., terpenoids, tannins, phenolic acids, flavonoids, and polyphenols) detected and recovered from other plant organs like stems, leaves, seeds, and their litter (Rice and Pancholy 1974; Coskun et al., 2017; Nardi et al., 2020; Lu et al., 2021; Wang et al., 2021). BNIs can suppress up to 90% of soil nitrification activity and increase soil N retention, providing plants with a competitive advantage for N fertilizer (Subbarao et al., 2009). However, relatively little is known about the exact mechanisms by which BNIs act on soil nitrifying microbes (Subbarao et al., 2015), with the vast majority of studies relying on screening tests with a single AOB *Nitrosomonas europaea* strain (Subbarao et al., 2006b; Zakir et al., 2008; Subbarao et al., 2013; Otaka et al., 2022). However, ammonia oxidation in soil is often dominated by ammonia-oxidising archaea (AOA) (Leininger et al., 2006), especially in acidic soils (Zhao et al., 2020) where BNIs release from plant roots is promoted (Di et al., 2018; Zhu et al., 2012; Zhang et al., 2019), and AOB *Nitrosospira* sp., rather than *Nitrosomonas*, in fertilized soils (Kowalchuk and Stephen, 2001; Norton et al., 2008). Hence, inhibition assays with a diverse range of soil-derived nitrifying strains could be a first and necessary step to define the spectrum of activity of NIs destined for use in agricultural settings (Papadopoulou et al., 2020). Such assays have been used for determining the activity of SNIs towards ammonia-oxidizing microbes (AOM) (Lehtovirta-Morley et al., 2013; Shen et al., 2013; Papadopoulou et al., 2020), and very recently of some BNIs (Kaur-Bhambra et al., 2022). These studies allowed for the estimation of inhibition thresholds based on the percentage reduction of ammonia oxidation (NO_3_^−^ accumulation) (Papadopoulou et al., 2020) or growth (μmax) (Kaur-Bhambra et al., 2022). Research on BNIs has been intensified in recent years, and new studies are reporting new plant-derived BNIs whose activity on the full breadth of soil nitrifying microbial groups remains unknown. This knowledge is required to get a first verification of their potential efficiency in agricultural settings, as well as their inhibition potential towards non-target soil microorganisms that might be expected based on the inherent role of some of these phytochemicals as plant defence agents (Dayan et al., 2010).

We aimed to determine the NI efficiency of multiple plant-derived molecules including (i) three root-exudated molecules with known BNI activity (MHPP, sakuranetin, and zeanone), (ii) four phytochemicals isolated from other than root, plant organs (caffeic acid, quinic acid, chlorogenic acid, and shikimic acid), and (iii) four analogues of statin (simvastatin), triazole (1-butyl-4-propyl-triazole, and 1,4-dibutyltriazole) and zeanone (2-methoxy-1,4-naphthoquinone), on liquid cultures of ecophysiologically and phylogenetically distinct soil-derived AOA and AOB strains. We expanded our assays to *Nitrobacter* nitrite-oxidizing bacteria (NOB), known as superior competitors than *Nitrospira* at high resources availability (e.g., fertilized soils) (Xia et al., 2011; Nowka et al., 2015, Papadopoulou et al., 2022), to gain insights on the impact of these compounds on other microbial players in the nitrification process that are known to be spatially and functionally connected with AOM (Jones and Hallin 2018). To standardize and interpret the effects observed, the stability of the selected NIs in the laboratory assays was also determined.

## 2. Materials and Methods

### 2.1. Microbial Strains and Growth Conditions

Our inhibition assays included two AOB strains, *Nitrosomonas europaea* ATCC 25978 and *Nitrosospira multiformis* ATCC 25196, two AOA strains, *Candidatus* Nitrosotalea sinensis Nd2 (Lehtovirta-Morley et al., 2014) and *Candidatus* Nitrosocosmicus franklandianus C13 (Lehtovirta-Morley et al., 2016a), and one NOB strain *Nitrobacter* sp. NHB1 (de Boer et al., 1991). All strains were grown aerobically in the dark without shaking. *N. multiformis* and *N. europaea* were grown at 28°C, in Skinner and Walker’s medium (SW) (Skinner and Walker, 1961) containing 1 mM NH_4_^+^ [(NH_4_)_2_SO_4_] and phenol red (0.5 mg L^−1^) as a pH indicator (pH 7.5-8.0). AOA *Ca.* N. franklandianus and *Ca.* N. sinensis, were incubated at 35°C in a medium supplemented with 1 mM and 0.5 mM NH_4_^+^ (NH_4_Cl), respectively. The former was cultured in HEPES-buffered modified freshwater medium (MFW) (pH 7.5; Lehtovirta-Morley et al., 2014), while the latter was grown in MES-buffered freshwater medium (FW) (pH 5.2; Lehtovirta-Morley et al., 2011). *Nitrobacter* sp. was grown at 28°C in FW medium (pH 5.2; Lehtovirta-Morley et al., 2011) supplemented with 0.5 mM NO_3_^−^ (NaNO).

### 2.2. BNIs and synthetic analogues

High purity (> 95%) analytical standards of sakuranetin, methyl 3-(4-hydroxyphenyl)-propionate (MHPP), 2,7-dimethoxy-1,4-naphthoquinone (zeanone), caffeic acid, quinic acid, chlorogenic acid, shikimic acid, simvastatin, 1-butyl-4-propyl-triazole, 1,4-dibutyltriazole, and 2-methoxy 1,4-naphthoquinone, were provided by Syngenta Crop Protection AG (Basel, Switzerland). Analytical standard of nitrapyrin (≥98%) was purchased from Sigma-Aldrich (Germany) and used as an internal control of the microbial cell response, according to a previous study by Papadopoulou et al. (2020). Detailed information on the tested compounds is provided in Table 1.

**Table 1.**
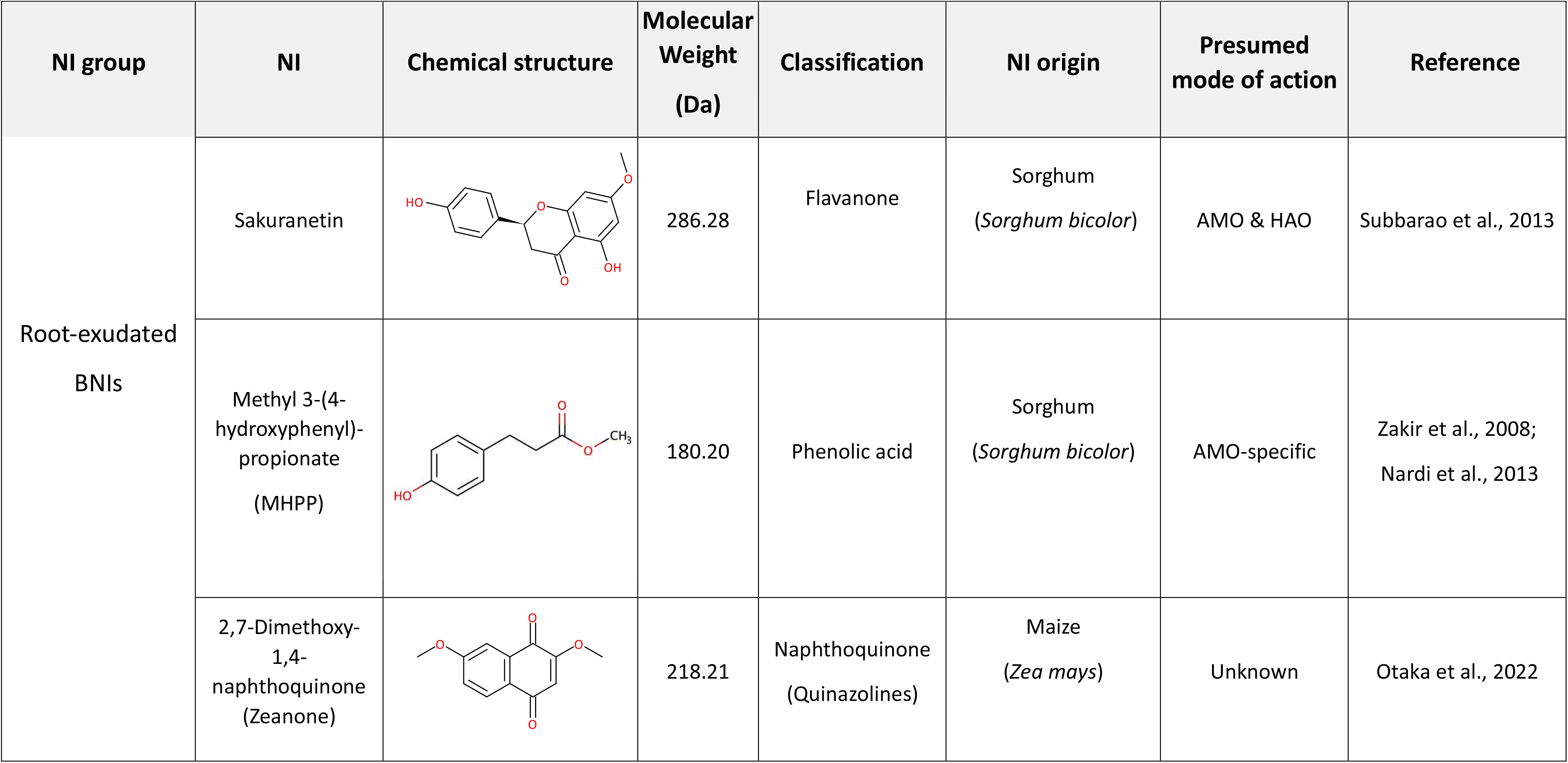
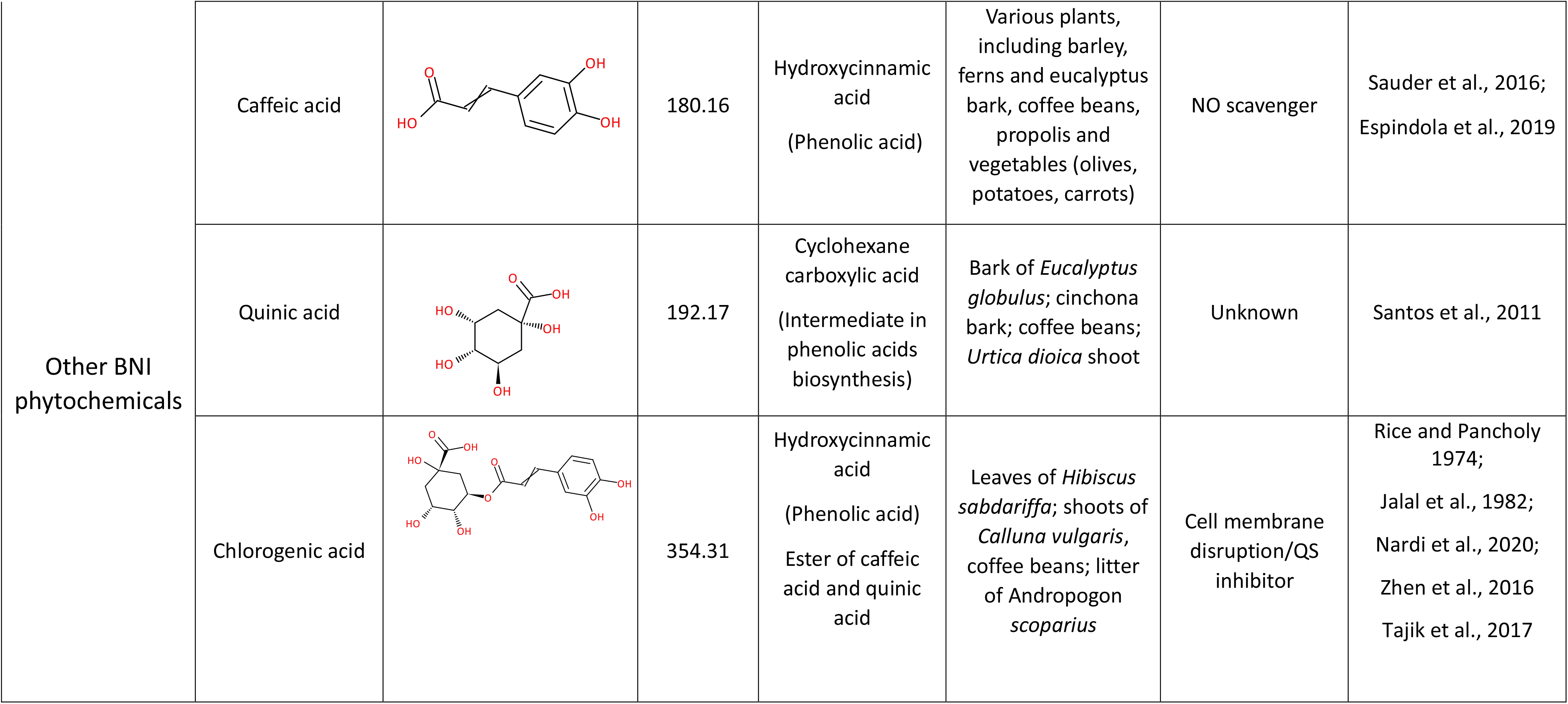
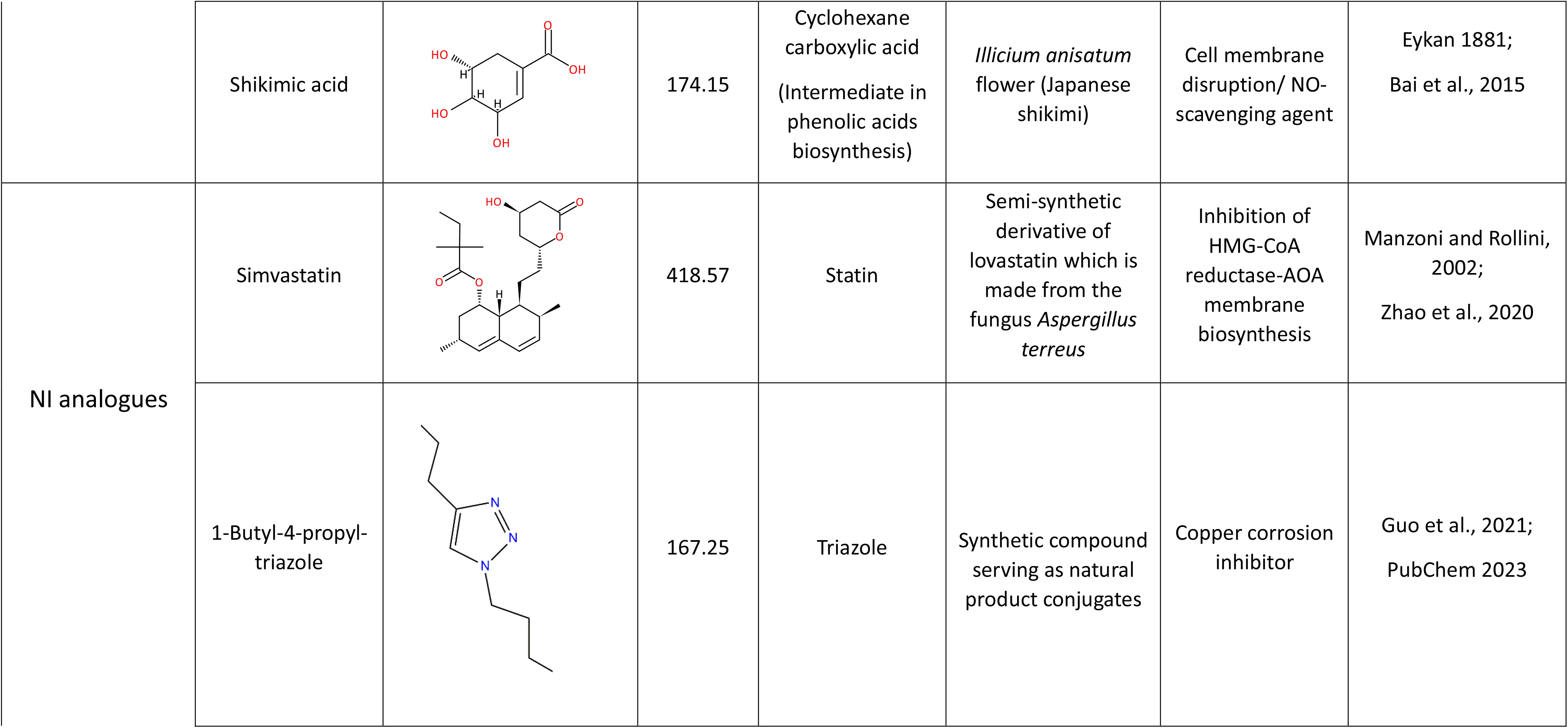
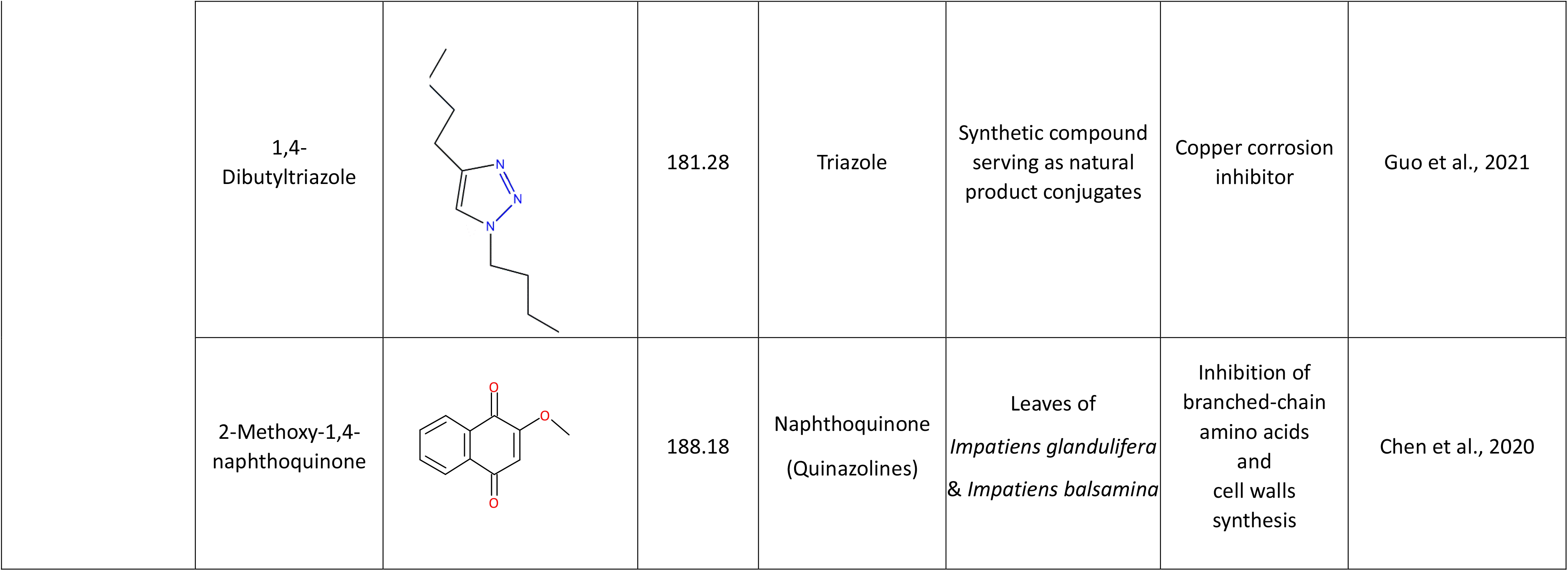
Summary of the nitrification inhibitors (NIs) used and detailed information about their (i) chemical structures; (ii) molecular weight (Da); (iii) chemical classification; (iv) origin; and (v) presumed mode of action.

### 2.3. Liquid Culture Inhibition Assays

To determine the inhibition thresholds per strain and compound, the activity of all NIs was tested in liquid batch cultures over a range of concentrations (see Table S1). Cultures were established in triplicate for each NI x concentration combination in 100-mL Duran bottles containing 30 mL of growth medium and inoculated with a 1% or 2% (vol/vol) transfer of exponentially growing culture of AOB or AOA/NOB respectively. All compounds except of quinic acid and shikimic acid were added to the cultures as filter sterilized dimethyl sulfoxide (DMSO) solutions due to their low water solubility. The final concentration of DMSO in all cultures was 0.1% (vol/vol), which did not exert a significant inhibitory effect to any of the isolates tested, in line with previous studies with the same isolates (Papadopoulou et al., 2020; Kaur-Bhambra et al., 2021). Quinic acid and shikimic acid were dissolved in sterile dH_2_O before addition in the cultures. All NIs were added to batch cultures at the beginning of the exponential growth phase. Triplicate cultures with the same inoculum amended with DMSO or sterile dH_2_O (in case of quinic acid and shikimic acid) without NIs were included as control. For each strain triplicate cultures were amended with a single concentration of nitrapyrin (5 μΜ, with known inhibitory activity to the tested strains based on Papadopoulou et al., 2020) to verify the consistency of our inhibition assays. For each of the tested NIs, triplicate non-inoculated cultures of SW (pH 7.8; Skinner and Walker, 1961) and FW medium (pH 5.2; Lehtovirta-Morley et al., 2011) were also prepared at one concentration level to determine the contribution of abiotic processes in the degradation of the tested compounds. It should be noted that these two media were selected to demonstrate the abiotic degradation of the NIs under alkaline and acidic conditions.

Upon inoculation all cultures were sampled at regular time intervals (once or twice daily) to determine the effect of NIs on the activity of the nitrifying microorganisms by measuring changes in nitrite concentrations. Nitrite concentrations were determined colorimetrically at 540 nm in a 96-well plate format assay by diazotizing and coupling with Griess reagent (Shinn, 1941). At specific cases (see Fig. S1) the effects of NIs on the growth of AOM and NOB were determined via q-PCR measurement of the abundance of the *amoA* and *nxrB* genes at selected time points based on the activity of the control treatment (early logarithmic and/or stationary phase). The degradation of NIs in the growth media was determined in samples collected at two time points (start: early logarithmic phase, end: early stationary phase) via HPLC.

### 2.4. q-PCR measurement of microbial growth of AOM and NOB

The abundance of *amoA* and *nxrB* genes was determined in a Biorad CFX Real–Time PCR system. DNA was extracted from a cell pellet obtained from 2-ml aliquots of the microbial cultures using the tissue DNA extraction kit (Macherey-Nagel, Germany). The *amoA* gene of *Ca.* N. sinensis was amplified with primers Arch-amoAF/Arch-amoAR (Francis et al., 2005), as described by Rousidou et al. (2013), and the *nxrB* gene of *Nitrobacter* sp. was quantified with primers nxrB-1F and nxrB-1R (Vanparys et al., 2007). The qPCR assays used the following thermal cycling conditions: 95°C for 3 min, followed by 40 cycles of 95°C for 30 s, 57°C for 20 s, 72°C for 30 s, with a final dissociation curve analysis. The abundance of *amoA* and *nxrB* was determined via external standard curves as described by Rousidou et al. (2013). qPCR amplification efficiency ranged from 87.9 to 90.9% with r^2^ values > 0.98.

### 2.5. Analysis of the stability of BNIs in liquid cultures

#### 2.5.1. Extraction methods

NIs were extracted from the growth media by mixing liquid culture with acetonitrile (MeCN) or methanol (MeOH) in a 1:2, 1:5, or 1:10 (v/v) ratio. The derived mixtures were vortexed for 1 min and stored at -20°C until HPLC analysis. Details of the extraction methods employed per compound is given in Supplemental material. Recovery tests at three concentration levels (in the range of the tested concentrations) showed recoveries of >70% for all compounds studied (Table S2). The sole exception was sakuranetin for which the achieved recoveries at the highest tested concentration (500 μΜ) were 28.2 and 33.7 % in the SW and FW medium, respectively.

#### 2.5.2. Chromatographic analyses

HPLC analyses was performed in a Shimadzu LC-20ADHPLC system equipped with an UV/VIS PDA detector. A Shimadzu GVP-ODs (4.6 mm by 150 mm, 5 mm) pre-column, connected to a RP Shimadzu VPODs (4.6 mm x 150 mm, 5 mm) column, was used for NIs separation. Chromatographic separation of all compounds was achieved at isocratic conditions as summarized in Table S3. Calibration curves obtained by the injection of standard solutions of NIs in MeOH or MeCN (in case of 2-methoxy-1,4-naphthoquinone and zeanone), ranging from 1 to 100 mg L^−1^ for caffeic acid and from 0.05 to 20 mg L^−1^ for all other compounds, were used for quantification.

### 2.6. Calculation of Inhibition Threshold Levels (EC_50_)

The concentration of each NI achieving 50% inhibition (EC_50_) in the activity of the AOB, AOA and NOB strains was calculated according to Papadopoulou et al. (2020). Briefly, dose-response modelling was performed using normalized data whereby nitrite concentration values were divided by the mean value of the matching control. Analyses were carried out using the dose response curves (drc) v3.0-1 package (Ritz and Streibig, 2005) of the R software.

### 2.7. Data analysis

Nitrite data were subjected to one-way analysis of variance (ANOVA), followed by Tukey’s post hoc test (p< 0.05). Variance between the EC_50_ values of the different NIs for one strain and between different strains for a given NI was analysed by one-way ANOVA, and Tukey’s post hoc test (P < 0.05). Comparison of mean EC_50_ values of AOB, AOA and NOB for all NIs was performed using the non-parametric Wilcoxon signed rank test.

## 3. Results

### 3.1. The inhibition of BNIs on the activity of AOM and NOB strains

#### 3.1.1. Root exudated BNIs

Sakuranetin induced a complete, non-reversible inhibition of the activity of *N. multiformis* at concentrations ≥ 250μΜ, but it did not significantly affect (p > 0.05) NO_3_^−^ production by *N. europaea* at the tested concentration range (10-500 μΜ). Conversely, the activity of both AOA strains was fully inhibited by sakuranetin at all tested concentrations (≥10 μΜ). NO_3_^−^ consumption by *Nitrobacter* sp. was fully inhibited at 250 μΜ, while a weaker inhibitory effect was observed at 500 μΜ (Fig. 1).

**Figure 1.**
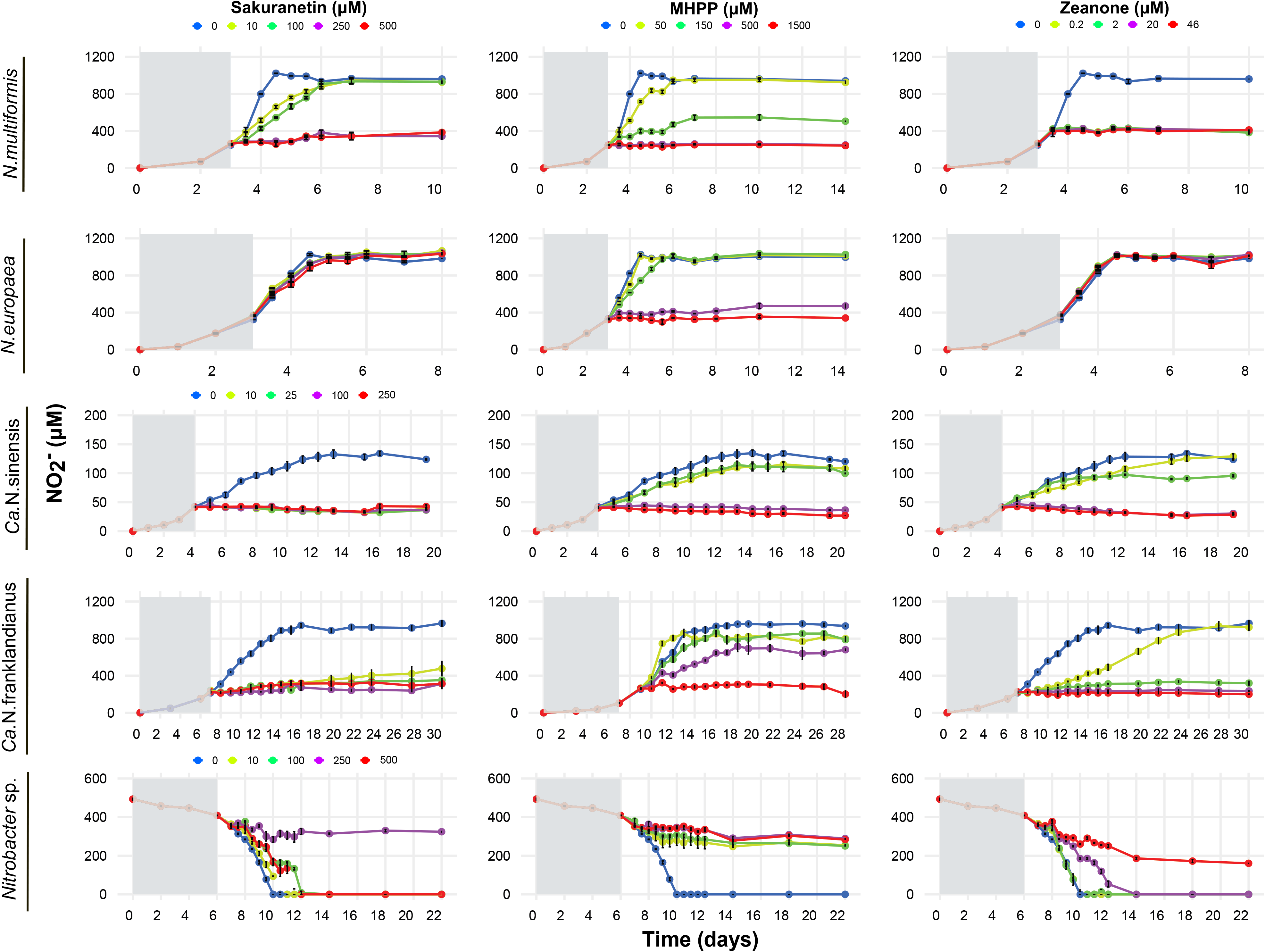
The effect of different concentrations of sakuranetin, MHPP, and zeanone, on the activity of the ammonia-oxidizing bacteria (AOB) *N. multiformis* and *N. europaea*, the ammonia-oxidizing archaea (AOA) *Ca.* N. sinensis and *Ca.* N. franklandianus, and the nitrite-oxidizing bacterium (NOB) *Nitrobacter* sp., determined by monitoring the production or consumption of nitrite. Error bars represent the standard error of the mean of triplicate cultures.

MHPP significantly reduced (p < 0.05) the activity of *N. multiformis* at levels ≥ 150 μΜ, and complete inhibition for both AOB strains was evident at ≥ 500 μΜ. Ammonia oxidation by *Ca.* N. sinensis and *Ca.* N. franklandianus was significantly suppressed (p < 0.05) at concentrations ≥ 500 μΜ. Finally, nitrite oxidation by *Nitrobacter* sp. was fully inhibited at all concentrations tested (≥ 50 μΜ) (Fig. 1).

Zeanone fully inhibited the activity of *N. multiformis* and of the two AOA strains at concentrations ≥ 0.2 μΜ and ≥2 μΜ, respectively, while no significant inhibition (p > 0.05) on the activity of *N. europaea* was observed at the tested concentration range (0.2-46 μΜ). Nitrite oxidation by *Nitrobacter* sp. was significantly (p<0.05) suppressed at 20 μΜ and completely halted at 40 μΜ of zeanone (Fig. 1).

#### 3.1.2. Phytochemicals

Caffeic acid, induced a complete, non-reversible inhibition of the activity of *N. multiformis* and *N. europaea* at concentrations ≥ 300 μΜ, while the same concentration of caffeic acid (300 μΜ) induced a temporal inhibitory effect in the activity of *Ca.* N. franklandianus (Fig. 2). On the other hand, *Ca.* N. sinensis was fully inhibited at lower concentrations (≥30 μΜ). *Nitrobacter* sp. was inhibited at concentrations ≤ 30 μΜ, whereas a rapid consumption of NO_3_^−^ was observed at the higher concentrations of caffeic acid ≥300 μΜ.

**Figure 2.**
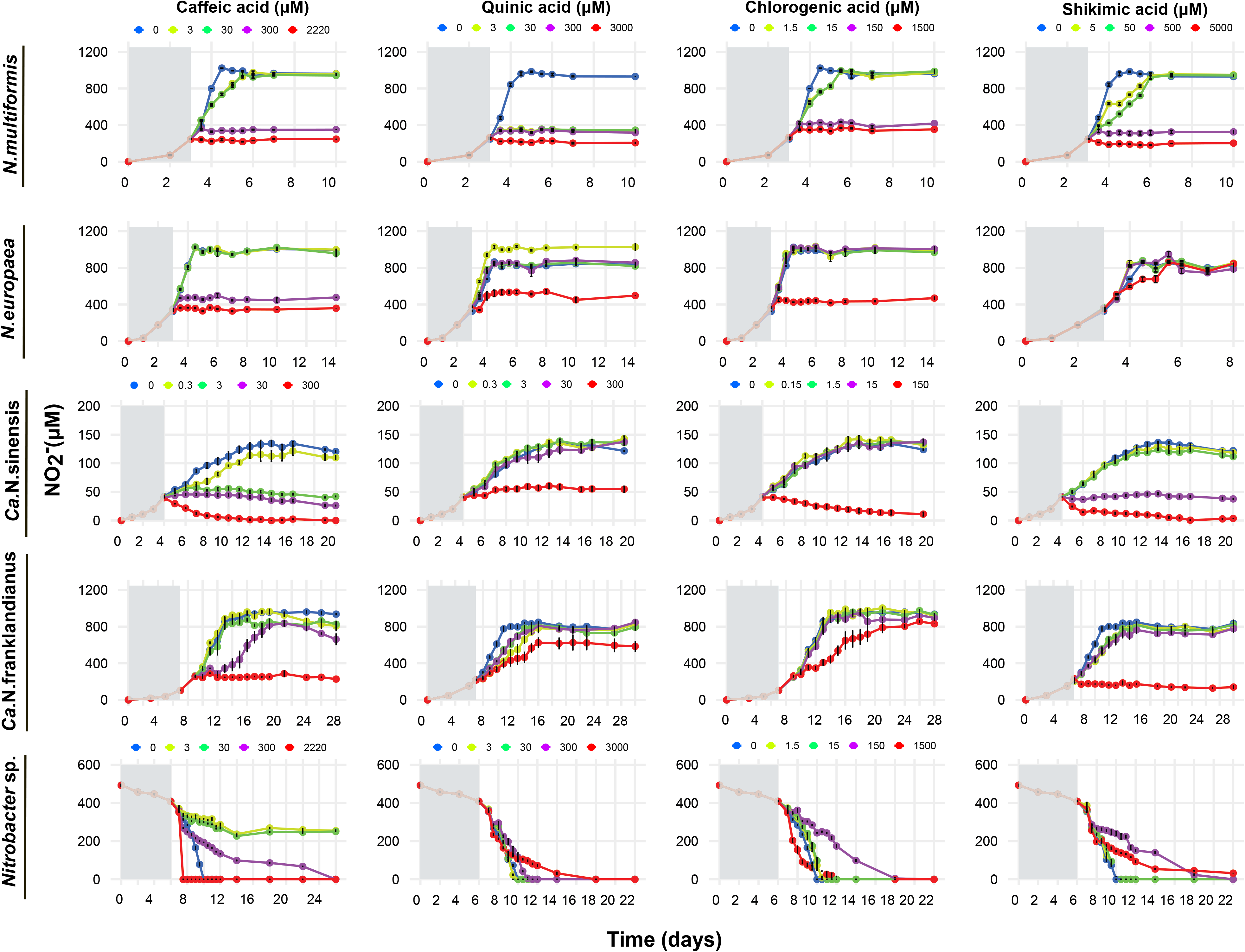
The effect of different concentrations of caffeic acid, quinic acid, chlorogenic acid, and shikimic acid on the activity of the ammonia-oxidizing bacteria (AOB) *N. multiformis* and *N. europaea*, the ammonia-oxidizing archaea (AOA) *Ca.* N. sinensis and *Ca.* N. franklandianus, and the nitrite-oxidizing bacterium (NOB) *Nitrobacter* sp., determined by monitoring the production or consumption of nitrite. Error bars represent the standard error of the mean of triplicate cultures.

Quinic acid fully inhibited ammonia oxidation by *N. multiformis* at concentrations ≥ 3 μΜ (Fig. 2). A complete, non-reversible inhibition of the activity of *N. europaea* and *Ca.* N. sinensis was observed only at the highest tested concentrations of 3000 μΜ. At the same concentration level quinic acid significantly reduced but not fully inhibited the activity of *Ca.* N. franklandianus. Finally, *Nitrobacter* sp. was not significantly affected (p > 0.05) by quinic acid at the tested concentration range (3-3000 μΜ).

Chlorogenic acid, induced a complete, non-reversible inhibition of the activity of *N. multiformis* and *N. europaea* at concentrations ≥ 150 μΜ and 1500 μΜ, respectively (Fig. 2). AOA activity was significantly reduced (p < 0.05) only at the concentration of 1500 μΜ with the inhibitory effect being non-reversible only in the case of *Ca.* N. sinensis (Fig. 2). Finally, we noted a significant inhibition of the nitrite oxidation activity of the *Nitrobacter* strain only at the 150 μΜ, while the higher tested level of chlorogenic acid (1500 μΜ) did not inhibit the activity of the NOB strain.

Shikimic acid significantly reduced the activity of *N. multiformis*, *Ca.* N. sinensis, and *Nitrobacter* sp. at concentrations ≥500 μΜ. In contrast, a complete, non-reversible inhibition of the activity of *Ca.* N. franklandianus was induced only at the highest tested concentration of 5000 μΜ, while the same concentration had no effect on *N. europaea* (Fig. 2).

#### 3.1.3. Analogues of NIs

Simvastatin did not significantly affect (p > 0.05) the activity of AOB and *Nitrobacter* sp., at the tested concentration range. In contrast, it significantly reduced (p < 0.05) the ammonia oxidation activity of *Ca.* N. sinensis and *Ca.* N. franklandianus at concentrations ≥ 5 μΜ and ≥ 15 μΜ, respectively (Fig. 3).

**Figure 3.**
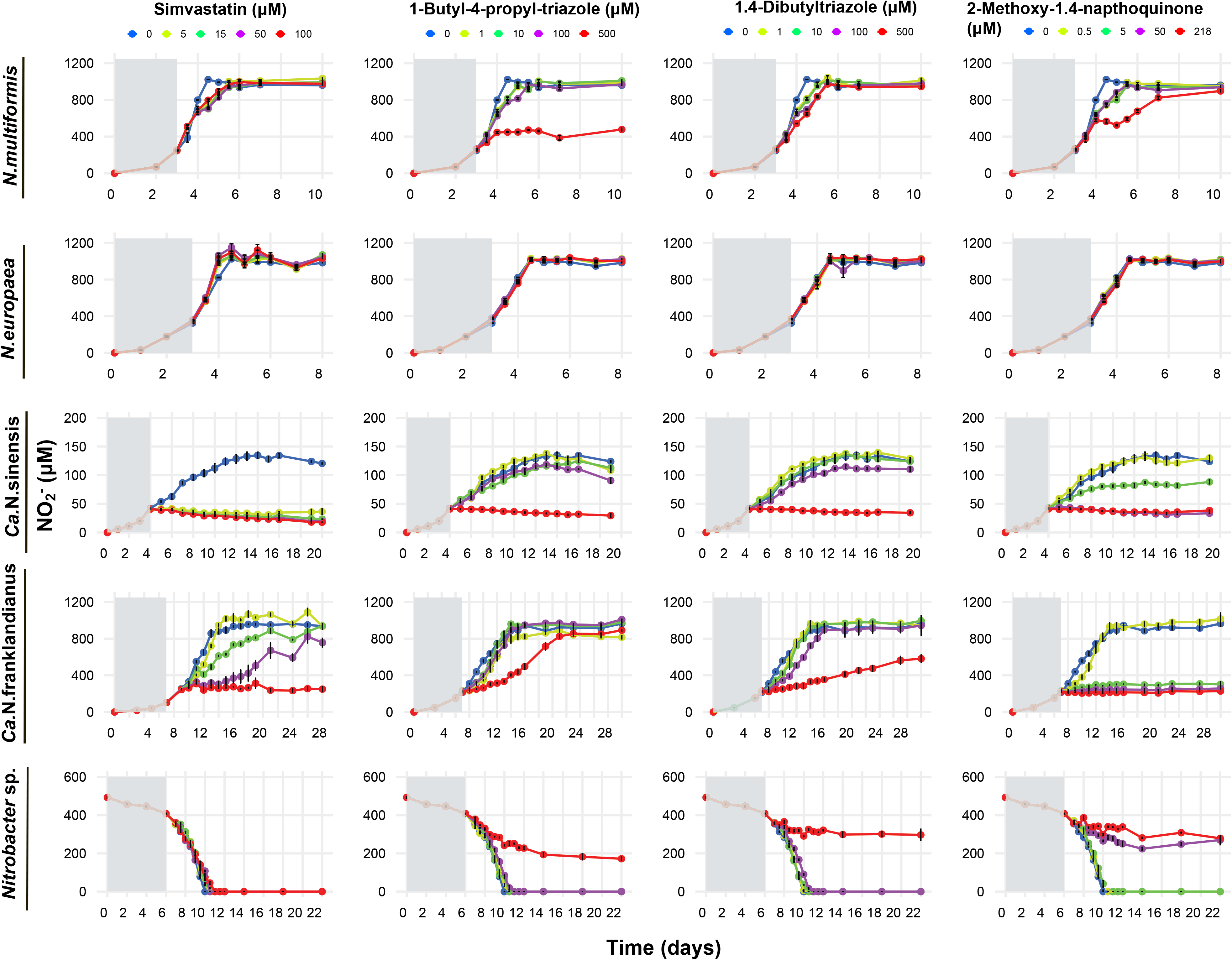
The effect of different concentrations of simvastatin, 1-butyl-4-propyl-triazole, and 1,4-dibutyltriazole, and 2-methoxy-1,4-naphthoquinone on the activity of the ammonia-oxidizing bacteria (AOB) *N. multiformis* and *N. europaea*, the ammonia-oxidizing archaea (AOA) *Ca.* N. sinensis and *Ca.* N. franklandianus, and the nitrite-oxidizing bacterium (NOB) *Nitrobacter* sp., determined by monitoring the production or consumption of nitrite. Error bars represent the standard error of the mean of triplicate cultures.

1-Butyl-4-propyl-triazole significantly inhibited (p <0.05) ammonia-oxidation by *N. multiformis*, AOA strains and *Nitrobacter* sp. only at the highest concentration tested, 500 μΜ, with the effect on *Ca.* N. franklandianus being temporal (Fig. 3). No effect on *N. europaea* was observed.

1,4-Dibutyltriazole did not significantly affect (p > 0.05) ammonia oxidation by AOB at the tested concentration range but induced a significant reduction on the activity of the AOA strains and of the *Nitrobacter* strain at the highest tested concentration of 500 μΜ (Fig. 3).

2-Methoxy-1,4-naphthoquinone only temporarily inhibited ammonia oxidation by *N*. *multiformis*, at the highest concentration tested, 218 μΜ, while at the same concentration level *N. europaea* activity was not significantly affected (p > 0.05). Ammonia oxidation by *Ca.* N. sinensis and *Ca.* N. franklandianus was significantly reduced at concentrations ≥5 μΜ (p<0.05). Nitrite consumption by *Nitrobacter* sp. was completely inhibited at concentrations ≥50 μM (Fig. 1).

#### 3.1.4. Nitrapyrin

Nitrapyrin, used as an internal control in our assays, at a single concentration (5 μΜ): (i) induced a complete, non-reversible inhibition of the activity of the AOB strains, (ii) imposed a significant inhibition of the activity of the AOA strains, with the effect on *Ca.* N. sinensis being temporal (iii) did not affect nitrite oxidation by the NOB *Nitrobacter* sp., in agreement with the data of Papadopoulou et al. (2020) (Fig. S3).

### 3.2. The effect of selected BNIs on the growth of AOM and NOB strains

During activity testing we noted a gradual decrease in the concentration of NO_3_^−^ in the cultures of *Ca.* N. sinensis amended with the highest concentrations of caffeic acid (300 μΜ), chlorogenic acid (150 μΜ), and shikimic acid (5000 μΜ) (Fig. 2). Although this could be attributed to a potential interaction under acidic conditions between these compounds and the produced NO_3_^−^, as reported before (Cotelle and Vezin, 2001; Napolitano and Ischia, 2002; Rabelo et al., 2015), we verified the inhibitory effect of these compounds by q-PCR monitoring of their growth. Indeed, the growth inhibition pattern of *Ca.* N. sinensis was congruent with NO_3_^−^ production patterns at concentrations of ≥ 3 μΜ (caffeic acid), 150 μΜ (chlorogenic acid), and ≥ 500 μΜ (shikimic acid) (Fig. S1).

Similarly, we monitored the growth of *Nitrobacter* sp., in the cultures amended with caffeic acid (300 and 2200 μΜ), chlorogenic acid (1500 μΜ), shikimic acid (500 and 5000 μΜ) and sakuranetin (500 μΜ) to verify the activity inhibition of *Nitrobacter* sp., since the NO_3_^−^ consumption patterns of *Nitrobacter* sp. deviated from a typical dose response pattern (Fig. 1 and 2). In contrast to the nitrite consumption measurements, all compounds imposed a significant reduction in the abundance of the *nxrB* gene of *Nitrobacter* sp., unlike sakuranetin where no significant inhibition of the growth of *Nitrobacter* sp. at the 500 μΜ level was observed (Fig. S2). We speculate that this inconsistency was due to the limited water solubility of sakuranetin, hence the relevant data from this concentration level were excluded from the calculation of the respective EC_50_ values for all nitrifying strains tested.

### 3.3. The persistence of NIs on AOM and NOB cultures

The persistence of sakuranetin, MHPP, and zeanone varied among the different cultures and NI concentration levels (Fig. S4). Caffeic acid and chlorogenic acid showed high degradation levels (*ca.* 80-100%) in AOB and *Ca.* N. franklandianus cultures, all growing in alkaline media, while lower degradation rates were observed in the *Ca.* N. sinensis and *Nitrobacter* sp. cultures (Fig. S5) both growing in acidic media. Shikimic acid, was generally stable and showed appreciable degradation only at the lower concentration levels of 5 and 50 μΜ (Fig. S5). Simvastatin was less stable under acidic growing conditions (Fig. S6). 1-Βutyl-4-propyl-triazole and 1,4-dibutyltriazole showed significant degradation levels only at the lowest concentration levels, while the persistence of 2-methoxy-1,4-napthoquinone varied in the different nitrifying cultures (Fig. S6). NIs showed similar degradation patterns in the inoculated and non-inoculated (abiotic controls) SW medium of *N. multiformis* (Fig. S4-S6). In contrast, MHPP, zeanone, simvastatin, and 2-methoxy-1,4-napthoquinone showed significantly higher degradation levels (p < 0.05) in the cultures inoculated with *N. europaea* compared to the corresponding non-inoculated SW medium. Similarly, sakuranetin, zeanone, chlorogenic acid, and simvastatin showed significantly higher degradation levels (p< 0.05) in the inoculated with *Ca.* N. sinensis FW medium compared to the respective non inoculated cultures (Fig. S4-S6). A more detailed description of the data regarding the persistence of NIs is provided in the Supplemental Material.

### 3.4. EC_50_ values of the tested NIs per strain

AOA (mean EC_50_ = 220.5 μM), were overly more sensitive than AOB (mean EC_50_ = 519.1 μM) to the NIs tested (p = 0.017), while *Nitrobacter* sp. (EC_50_ = 390.6 μM) showed not significant difference in its sensitivity compared to AOB and AOA (p > 0.05) (Fig. 4).

**Figure 4.**
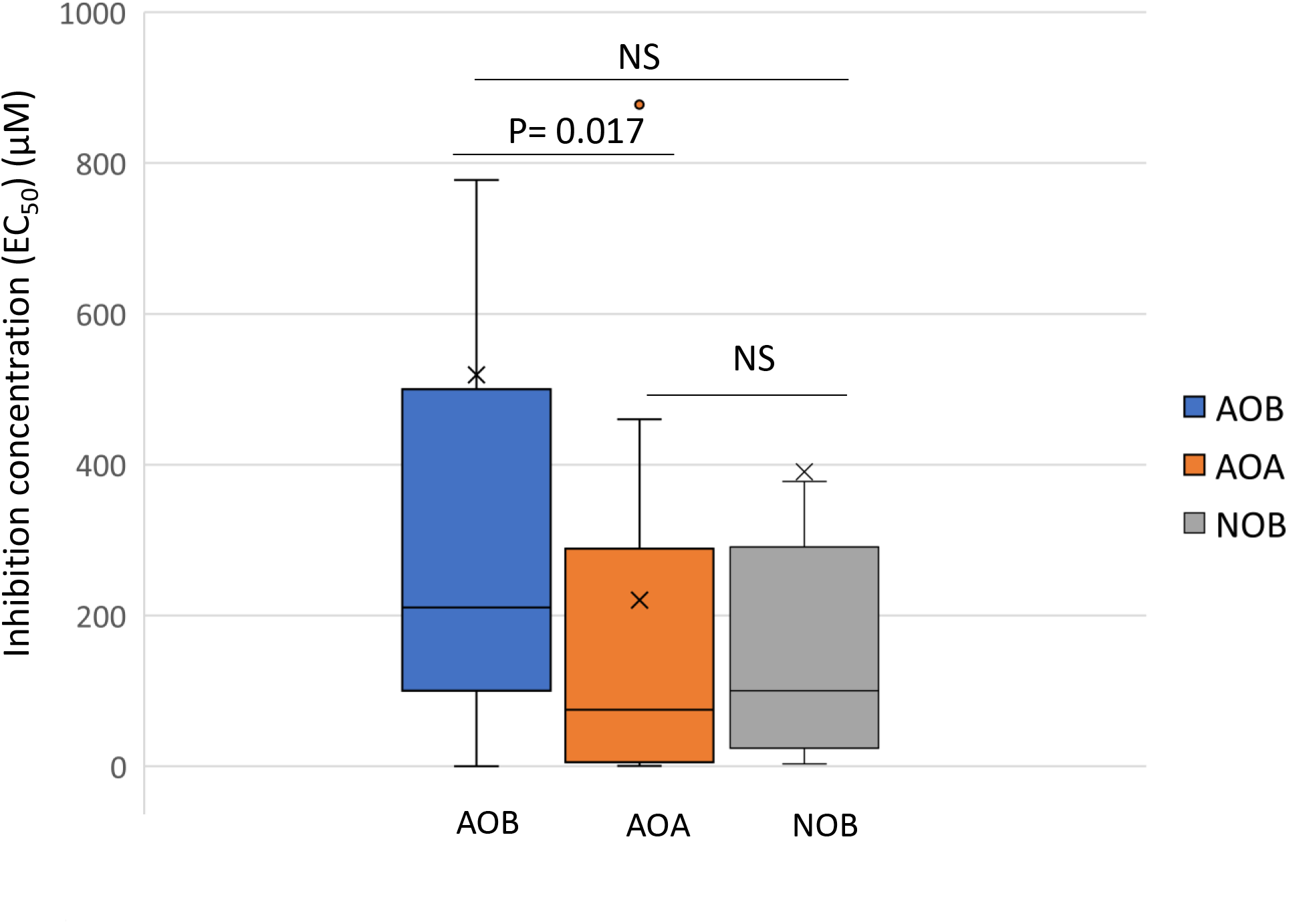
The mean EC_50_ values (μΜ) derived from all tested nitrification inhibitors (NIs) for ammonia-oxidizing bacteria (AOB), ammonia-oxidizing archaea (AOA) and the nitrite-oxidizing bacterium (NΟΒ). NS, denotes not significant differences (p > 0.05). In cases where no EC_50_ could be descent from the statistical analysis the lowest or the highest tested concentrations were used instead.

The two tested AOB strains, *N. multiformis* and *N. europaea,* varied in their sensitivity to the tested NIs with the former being consistently more sensitive (p <0.05) than the later to eight out of the 11 NIs tested (Fig. 5, Table S4). Zeanone (EC_50_ <0.2 μM) and quinic acid (EC_50_ <3 μM) were the most potent inhibitors of *N. multiformis*, while caffeic acid (EC_50_ =203.3 ± 15.8 μM) and MHPP (EC_50_ =339.3 ± 32.4 μM) were the most potent inhibitors of *N. europaea* (Fig. 5). The most potent NIs towards AOA were zeanone, its analogue 2-methoxy-1,4-naphthoquinone, caffeic acid, sakuranetin and simvastatin (Fig. 5). *Ca.* N. sinensis was generally more sensitive than *Ca.* N. franklandianus to six out of the 11 NIs tested. Zeanone and its analogue 2-methoxy-1,4-naphthoquinone were significantly more potent to *Ca.* N. franklandianus compared to *Ca.* N. sinensis, while 1-butyl-4-propyl-triazole and 1,4-dibutyltriazole were equally potent to both AOA strains (Fig. 5, Table S5). Regarding the NOB strain *Nitrobacter* sp., caffeic acid (EC_50_ <3 μM) was its most potent inhibitor. This was followed by MHPP, zeanone, 2-methoxy-1.4-naphthoquinone, and chlorogenic acid which were equally potent against the NOB strain (p > 0.05). Amongst them MHPP showed the lowest (EC_50_ =7.4 ± 1.3 μM) and chlorogenic acid the highest EC_50_ value (71.9±8.9μM) (Fig. 5).

**Figure 5.**
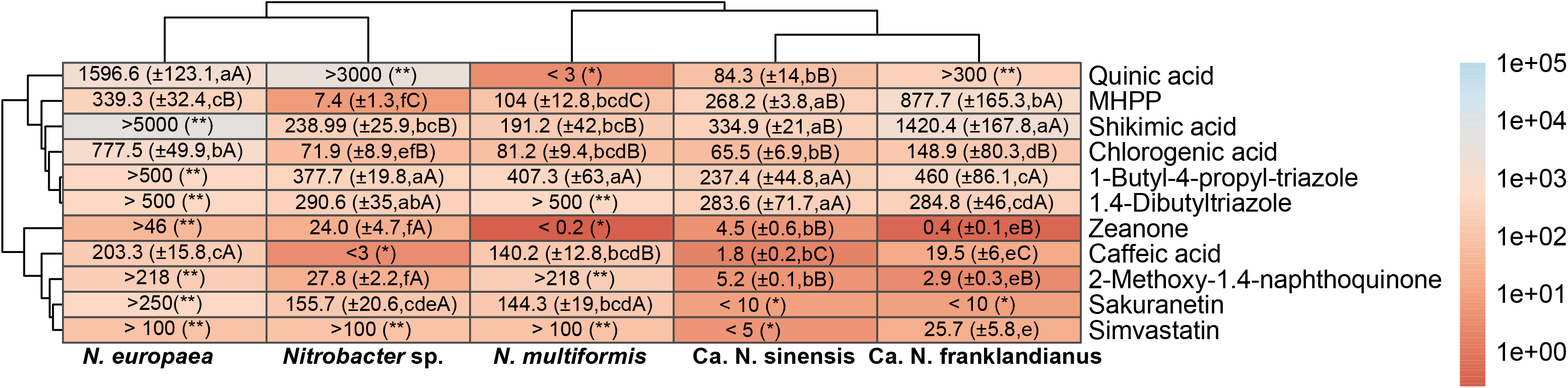
Mean EC_50_ values (μM) of the tested nitrification inhibitors (NIs) calculated based on their inhibitory activity on the ammonia or nitrite oxidation capacity of the tested ammonia-oxidizing bacteria (AOB), ammonia-oxidizing archaea (AOA), and the nitrite-oxidizing bacterium (NOB). Standard errors of the mean values (denoted by±) are given in brackets. Upper case letters indicate significant differences (p < 0.05) between microorganisms for each individual NI, and lower-case letters indicate significant differences (p < 0.05) between NIs for each tested microorganism. The asterisk(s) denotes that no EC_50_ could be descent from the data. One asterisk (*) indicates that EC_50_ value is lower than the minimum tested concentration, while two asterisks (**) indicate that the EC_50_ value is higher than the maximum tested concentration. Dendrograms based on the Euclidean distances and the complete linkage clustering method using log transformed mean EC_50_ values are presented for identifying NIs. The table is color-coded by orders of magnitude for EC_50_ values according to the colour legend.

## 4. Discussion

We determined the inhibitory activity of a wide range of plant-derived molecules categorized as (i) root-derived BNIs (ii) phytochemicals with potential BNI activity and (iii) analogues of statins, triazoles and zeanone, on a range of AOA and AOB strains. The first group was composed of known BNIs like sakuranetin, MHPP and zeanone. Sakuranetin exhibited higher inhibitory activity towards AOA isolates as also observed by Kaur-Bhambra et al. (2021). Regarding *N. europaea* and *Ca.* N. franklandianus, Kaur-Bhambra et al. (2021) reported an IC_80_ of 259 μM and 173 μM, respectively, compared to our EC_50_ values of > 250 μM and < 10 μM, for the same AOB. The differences in the inhibition thresholds between the two studies could be attributed either to the different *N. europaea* strains tested (ATCC 19718 instead of ATCC 25978 in our study) or most probably to the different approaches used for the calculation of the inhibition thresholds (concentration of each BNI leading to 80% inhibition of μmax vs concentration leading to 50% inhibition of nitrite production in our study). MHPP was the only tested BNI that showed higher potency towards AOB suppressing their activity at concentrations ≥ 150 μΜ compared to ≥ 500 μΜ for AOA isolates. This is in line with previous studies which showed 80% inhibition of AOB (*N. europaea*, *N. multiformis*, *N. tenuis*, and *N. briensis*) and AOA (*Ca.* N. sinensis, *Ca.* N. franklandianus, and *N. viennensis*) growth at levels varying from 80 to 545 μΜ for AOB and from 149 to 1028 μΜ for AOA (Kaur-Bhambra et al., 2021). In accord with our findings for MHPP inhibition on *N. europaea* (EC_50_ 339.3 ±32.4), Lu et al. (2021) using a different *N. europaea* stain (NBRC 14,298) reported a similar ED_50_ value of 295 μM. Zeanone, a novel naphthoquinone BNI isolated from maize root exudates (Otaka et al., 2022), was reported with an ED_80_ of 8 μM, determined using the recombinant *N. europaea* assay. Based on our study, which provides a broader characterization of its inhibitory activity, zeanone was a strong inhibitor of *N. multiformis* (EC_50_ < 0.2 μM) and of the two AOA isolates (EC_50_ = 4.5±0.6 μM and 0.4±0.1 μM for *Ca.* N. sinensis and *Ca.* N. franklandianus, respectively) but failed to inhibit *N. europaea* in the tested concentration range.

The second group of compounds tested included phytochemicals with potential NI activity like caffeic, quinic, chlorogenic and shikimic acid. As it was expected by its reported mode of action, as NO scavenger (Sueishi et al., 2011), caffeic acid showed a higher inhibition activity on AOA (mean EC_50_ = 10.7 μΜ) compared to AOB strains (mean EC_50_ = 171.7 μΜ). AOA are known to be inhibited by NO scavengers like PTIO (2-phenyl-4,4,5,5-tetramethylimidazoline-1-oxyl-3-oxide) which has been interpreted as evidence that NO is an intermediate in the AOA pathway, but not in AOB (Martens-Habbena et al. 2015; Kozlowski et al. 2016a). This hypothesis was cancelled by Caranto and Lancaster (2017) who showed that NO is also an intermediate in ammonia oxidation by AOB which are also inhibited by NO-scavengers, albeit at higher concentrations than AOA. The NO scavenging activity of caffeic acid has been previously tested against three AOA strains (*N. maritimus* SCM1, AOA-6f and AOA-G6), and the AOB *N. europaea* strain ATCC 19718 by Sauder et al. (2016) who reported an EC_50_ < 10 μM for *N. maritimus* and AOA-6f, and an EC_50_ of 84.3 μM for AOA-G6, in line with our study. However, the same authors did not observe any inhibition of *N. europaea* by caffeic acid (EC_50_ of > 300 μM) compared to our findings where an EC_50_ = 203.3 μM was calculated, using a different *N. europaea* strain. The NI activity of quinic acid was assessed for the first time and showed variable inhibition levels being more inhibitory towards *N. multiformis* (EC_50_ < 3 μΜ) and *Ca.* N. sinensis (EC_50_ 84.3±14.0 μΜ) compared to *N. europaea* (EC_50_ 1596.6 ±123.1 μΜ), and *Ca.* N. franklandianus (EC_50_ >300 μΜ). Chlorogenic acid, an ester of caffeic acid and quinic acid, showed an intermediate inhibitory activity between these two compounds, suggesting a potential influence of chemical structure to their inhibition mechanism. Earlier studies showed that both caffeic acid and chlorogenic acid could inhibit nitrification at a concentration range of 0.1 to 1000 μΜ (Rice and Pancholy 1973), in line with our study. Shikimic acid completely inhibited the activity of *N. multiformis* and *Ca.* N. sinensis, at lower concentrations (≥500 μΜ) than these found to suppress *Ca.* N. franklandianus (complete inhibition at 5000 μΜ). At the same concentration range (up to 5000 μΜ) *N. europaea* was only slightly inhibited, in line with previous studies that reported no inhibitory effect of shikimic acid on *N. europaea* ATCC 19718 strain at concentrations up to 2500 μM (Sun et al., 2016; Lu et al., 2021).

The third group of potential NIs, included analogues of compounds with known inhibitory activity towards AOM like simvastatin (analogue of lovastatin) (Manzoni and Rollini, 2002), 1-butyl-4-propyl-triazole and 1,4-dibutyltriazole (triazole analogues) and 2-methoxy-1,4-naphthoquinone (analogue of zeanone). Simvastatin selectively inhibited both AOA isolates but did not affect the AOB strains, in accord with Zhao et al. (2020). The selectivity of simvastatin towards AOA is due to its known activity as an inhibitor of 3-hydroxy-3-methylglutaryl-CoA (HMG-CoA) reductase, a crucial enzyme in the mevalonate pathway in archaeal cell membrane biosynthesis (Lam and Doolittle, 1992; Miller and Wolin, 2001), with AOB using an alternative pathway (Jain et al., 2014). The two triazole derivatives selectively inhibited AOA strains, showing a much weaker effect on AOB. Although there are no data available regarding the effect of these compounds on pure cultures of soil nitrifiers, previous studies demonstrated their high efficacy in inhibiting nitrification in soil (Dixit et al., 2021; McCarty and Bremner 1989), and highlighted their potential use as NIs that can be synthesized with good yields and high atom economy from readily available starting materials (Taggert et al., 2021). Finally, 2-methoxy-1,4-naphthoquinone showed a distinct higher inhibition potential for AOA vs AOB, similarly to its chemical analogue zeanone, although the latter was clearly more inhibitory to *N. multiformis* compared to 2-methoxy-1,4-naphthoquinone which showed no activity to AOB isolates.

When the data from all 11 tested compounds and AOM strains were looked comparatively between and across AOM groups we identified clear trends of selectivity in the inhibition between AOA and AOB as well as within the different groups of AOM tested. As a first general hint we noted a higher sensitivity of AOA compared to AOB in 10 out of the 11 potential BNI compounds tested with the sole exception of MHPP. Previous culture-based studies on the sensitivity of AOM to SNIs and BNIs have shown a rather selective inhibition of AOB by SNIs (Papadopoulou et al., 2020; Shen et al. 2013; Taylor et al. 2013) and a stronger inhibition of AOA by BNIs (Kaur-Bhambra et al. 2022). The variable sensitivity of soil AOM to different NIs may arise from fundamental differences in their biochemistry and physiology (Hodgskiss et al., 2023; Schleper and Nicol, 2010) and could be related to the particular mode of action of each NI at cellular level with some of them (e.g., simvastatin) targeting group-specific (AOA) biochemical pathways.

We further observed differences in the sensitivity between the AOA and the AOB strains tested. *N. multiformis* was consistently more sensitive than *N. europaea*, while the sensitivity of AOA strains was compound-depended, with *Ca.* N. franklandianus being less sensitive than *Ca.* N. sinensis to six out of the 11 NIs tested. Variations in the sensitivity of different AOA and AOB strains to SNIs (Papadopoulou et al., 2020; Wright et al. 2020; Zhao et al. 2020) and BNIs (Kaur-Bhambra et al., 2021) has been reported before. Papadopoulou et al. (2020) observed a higher sensitivity of *N. multiformis* over *N. europaea* to the potent SNI quinone imine and of *Ca.* N. sinensis over *Ca.* N. franklandianus to the commercial SNIs DCD and DMPP. Kaur-Bhambra et al. (2021) demonstrated the higher sensitivity of *Ca.* N. sinensis over *Ca.* N. franklandianus (and *Nitrososphaera viennensis*) and of *N. multiformis* over *N. europaea* to various root- and shoot-derived BNI compounds. The reasons for the different sensitivities of the tested AOB and AOA strains to the potential BNIs studied is most probably driven by differences in their metabolism, ecophysiology, and other cellular characteristics. For example, the membranes of all AOA studied contain crenarchaeol, a glycerol dialkyl glycerol tetraether lipid produced exclusively in archaea of the phylum Thaumarchaeota (Elling et al., 2017; Schouten et al., 2013). However, the proportion of crenarchaeol was found to be much higher in all tested neutrophilic AOA (including *Ca.* N. franklandianus) compared to the acidophilic *Ca.* Nitrosotalea strains (Lehtovirta-Morley et al., 2016b). This could potentially explain the lower sensitivity of *Ca.* N. frankandianus to NIs like simvastatin, shikimic acid and chlorogenic acid presumed to interfere with membrane biosynthesis (Zhao et al., 2020) or affect its integrity (Bai et al., 2015; Nardi et al., 2020). The higher tolerance of *Ca.* N. franklandianus to NIs has been also presumed to be associated with its capacity to produce extracellular polymeric substances (EPS) leading to aggregate formation that could potentially prevent NIs, and especially hydrophilic compounds like quinic acid and shikimic acid, from accessing the surface of cells attached to the hydrophobic EPS matrix (Papadopoulou et al., 2020; Gao et al., 2007). This production of EPS is a common feature for all *Ca.* Nitrosocosmicus isolates (Jung et al., 2016; Lehtovirta-Morley et al., 2016a; Sauder et al., 2017; Alves et al., 2019; Liu et al., 2019) and has been also reported as an AOB defence mechanism against NIs (Powell and Prosser, 1991). Regarding AOB, *N. europaea* and *N. multiformis* possess a largely different set of oxidative stress response proteins, alkyl hyperoxide reductase vs superoxide dismutase and rubrerythrin, respectively (Zorz et al., 2018), that might exhibit different efficiencies to oxidative stress imposed by certain BNIs. In addition, *N. europaea* possess a cytochrome P460 (CytL) that is involved in the oxidation of NH_2_OH and NO to NO_3_^−^, and the NO-responsive transcriptional regulators NsrR and NnrS known to regulate the expression of multiple genes involved in nitrosative stress responses. Whereas all this enzymatic arsenal against nitrosative stress, potentially imposed by BNIs, is not present in *N. multiformis* (Kozlowski et al., 2016b). Alternatively, the wider range of membrane protein transporters being operative in *N. europaea* have been suggested as a potential mechanism to cope with the inhibitory effects of chemicals like NIs (Zorz et al., 2018).

We expanded our tests to a *Nitrobacter* sp., as a representative of NOB, whose response to BNIs remains unknown. NOB are fuelled by substrates produced by AOM, and their activity results in the rapid conversion of potentially hazardous (for both plants and microbes) NO_3_^−^ to NO_3_^−^ that it is consumed by plants and soil microbes (Daims et al., 2016). In heavily fertilized agricultural soils, the NOB-derived NO_3_^−^ contributes to N losses and environmental pollution through NO_3_^−^leaching and subsequent production of nitrogen oxides (e.g., N_2_O) through the denitrification process (Raun and Johnson, 1999). We demonstrated that caffeic acid, MHPP, zeanone, 2-methoxy-1.4-naphthoquinone, and chlorogenic acid, were suppressive to *Nitrobacter* sp. at concentrations < 100 μΜ. Amongst them, caffeic acid and MHPP were the two NIs that showed higher activity to the *Nitrobacter* strain compared to the AOA and AOB strains tested. The impact of these compounds on the activity of both AOM and NOB could significantly impact the direction and degree of nitrogen transformation during nitrification, and especially in cases where NIs inhibit NOB to a greater extent than AOM. This could potentially lead to nitrite accumulation in soil, raising serious concerns for environmental quality (e.g., increased nitrite driven N_2_O production) and agricultural production (e.g., nitrite toxicity to plants). We provide pioneering data regarding the impact of BNIs and their synthetic analogues on a pure NOB culture. Further studies extended to other NOB, including the widely distributed and diverse *Nitrospira* species, are required to determine the full inhibitory potential of BNIs on soil NOB.

To date the overwhelming majority of BNI activity testing has been performed with a single genetically modified AOB *Nitrosomonas europaea* strain (Subbarao et al., 2006b; Zakir et al., 2008; Subbarao et al., 2013; Otaka et al., 2022). This approach was initially developed to monitor the nitrification process in wastewater treatment plants (Iizumi et al., 1998), where *N. europaea* represents a key ammonia-oxidizing microbe, and it was then adopted for screening root exudates for BNI activity (Subbarao et al., 2006b; Zakir et al., 2008; Subbarao et al., 2013; Otaka et al., 2022). Kaur-Bhambra et al. (2021) assessed the activity of BNIs produced by plant roots and shoots on selected AOB and AOA strains and questioned the relevance of *N. europaea* as a single indicator of BNI activity based on its highly variable response to BNIs compared to strains that are more representative of natural AOM soil assemblages. Our study verified the high variation in the sensitivity of *N. europaea* to BNIs, compared to *N. multiformis* which constitutes a more representative soil AOB, reinforcing the need for NI assays with diverse strains of soil representative AOM. In addition, inhibition thresholds of BNIs (i.e., sakuranetin, MHPP, zeanone, and caffeic acid) estimated for *N. europaea* in this study differ significantly from those reported by other researchers using either a wild-type (Kaur-Bhambra et al., 2021; Sauder et al., 2016; Sun et al., 2016;) or a genetically modified *N. europaea* strain (Subbarao et al., 2006b; Zakir et al., 2008; Subbarao et al., 2013; Otaka et al., 2022) and different experimental approaches (e.g., detection of inhibition based on measurements of μmax, NO_3_^−^ production or changes in fluorescence, duration of bioassays from 30 min to several days; determination of IC_80,_ IC_70_, or EC_50_ values). All these might imply an inherent variability in the response of *N. europaea* to in vitro testing conditions further stressing its unsuitability for use as a sole indicator of NI activity in screening assays. In addition, our findings highlight the need for standardisation of the existing and under development high-throughput BNIs screening biological systems to obtain more meaningful and comparable information.

Finally, an important limitation of the currently available BNI screening systems is the lack of data regarding the persistence and stability of these chemicals under the testing conditions. These are often essential for the interpretation of the inhibitory effects observed, directly associated with the duration of the exposure to the tested compounds. We provide first results on the stability and persistence of BNI compounds during laboratory incubation. We noted that some of the tested chemicals showed a pH-dependent stability (simvastatin), while others were overly unstable (caffeic and chlorogenic acid) or persistent (triazole derivatives) in all growth media. It is worth noting the potential involvement of *N. europaea* and *Ca*. N. sinensis in the cometabolic biodegradation of certain NIs (e.g., zeanone and simvastatin) serving as non-growth substrates for AOM strains in the presence of NH_3_/NH_4_^+^ (growth substate). The potential catabolism of NIs by AOM is not unexpected considering earlier evidence that the AMO enzymes of both AOB (e.g., *N. europaea*) and AOA (e.g., *N. gargensis)* could transform co-metabolically a wide range of aliphatic and aromatic substrates (Hyman et al., 1988; Rasche et al., 1991; Keener and Arp, 1994; Chang et al., 2003) including pharmaceuticals/antibiotics (e.g., β-lactams, fluoroquinolones, and sulphonamides) through distinct biodegradation pathways involving deamination, hydroxylation, and co-oxidation (Li et al., 2022; Zhou et al., 2019). Still the involvement of the tested AOM strains in the degradation of the tested NIs is a particularly interesting aspect which warrants further investigation.

## 5. Conclusions

We provided a systematic assessment of the inhibition potential of multiple BNI molecules on the activity of soil derived AOB (*Nitrosospira multiformis* and *Nitrosomonas europaea*), AOA *(Ca.* Nitrosotalea sinensis and *Ca.* Nitrosocosmicus franklandianus), and NOB (*Nitrobacter* sp.) strains, which represent main nitrifier lineages in soil. Accurate analytical methods for determining the degradation and stability of the NIs in the liquid cultures of the nitrifying isolates were developed and validated, in parallel. Our study (i) offers benchmarking knowledge of the activity of diverse BNIs and analogues to a range of soil AOM and *Nitrobacter* NOB, whose response to BNIs was unknown, (ii) provides unprecedented data on the inhibition potential of the recently discovered hydrophobic maize isolated BNI, zeanone, and its natural analogue, 2-methoxy-1,4-naphthoquinone, on nitrification; (iii) verifies the selective activity of the currently available BNIs on AOA and stresses the necessity for elucidation of the inhibition mechanisms against target AOM, (iv) illustrates the variation in the sensitivity of different AOM isolates to BNIs. This further highlights the limitations of studies relying on a single AOB strain and argues for testing against a broad range of ecophysiologicaly and phylogenetically distinct soil-derived nitrifiers to extrapolate a robust estimate of the NI potency of plant derived compounds; (v) provides first evidence for the stability of BNIs during biological screening allowing for accurate and comparable determination of inhibition thresholds; and (vii) demonstrates the potential inhibitory activity of BNIs on nitrifying microbial groups beyond AOM, modulating downstream processes in N cycling like NOB. The potency of the BNIs tested on *Nitrobacter* sp., questions the specificity of certain BNI molecules towards AOM that, like other phytochemicals, might have a wider antimicrobial activity, beyond just prokaryotic nitrifiers, that should be assessed before further use in agricultural settings.

## Acknowledgments

This project has received funding from Syngenta Crop Protection AG, Basel, Switzerland.

## Notes

### Competing Interest Statement

The authors have declared no competing interest.

