## Supplemental Material for "Assessing the activity of different plant-derived molecules and potential biological nitrification inhibitors on a range of soil ammonia- and nitrite- oxidizing strains"

### Material and Methods

#### Nitrification inhibitors (NIs) extraction from liquid cultures

For the extraction of sakuranetin and chlorogenic acid residues 0.1 mL of the liquid culture was mixed with 0.4 mL of methanol (1:5 vol/vol). MHPP and shikimic acid residues were extracted from all growth media by mixing 0.05 mL of liquid culture with 0.45 mL of methanol (1:10 vol/vol). Zeanone and 2-methoxy 1,4-naphthoquinone were extracted by mixing 0.25 mL liquid culture with 0.25 mL of acetonitrile (1:2 vol/vol). Caffeic acid was extracted from SW and MFW growth medium by mixing 0.05 mL of liquid culture with 0.45 mL of methanol (1:10 vol/vol) and from FW growth medium by mixing 0.1 mL of liquid culture with 0.4 mL of methanol (1:5 vol/vol). Simvastatin residues were extracted from SW and MFW medium by mixing 0.25 mL liquid culture with 0.25 mL of methanol (1:2 vol/vol), and from FW medium by mixing 0.05 mL of liquid culture with 0.45 mL of methanol (1:10 vol/vol). 1-Butyl-4-propyl-triazole, and 1,4-dibutyltriazole residues were extracted by mixing 0.25 mL liquid culture with 0.25 mL of methanol. Nitrapyrin residues were extracted from all growth substrates by mixing 0.3 mL liquid culture with 0.7 mL of acetonitrile, according to Papadopoulou et al. (2020). The derived mixtures were vortexed for 1 min and stored at -20°C until analysis. Quinic acid residues were not determined due to the inability to detect this compound in the available HPLC-DAD system.

#### Recovery tests and validation of the extraction methods

Analysis of fortified samples of the different growth media was conducted to verify the extraction efficiency of the developed methods (as described above for the different nitrifying cultures). For this purpose, two growth media (i) SW (pH 7.8; Skinner and Walker, 1961) and (ii) FW (pH 5.2; Lehtovirta-Morley et al., 2011), differing in chemical composition and pH, were used. Triplicate samples (5 mL) of each growth medium were spiked with the DMSO (50μL; 0.1% vol/vol) or water (0.883 mL, 17.7% vol/vol) solutions of the nitrification inhibitors (NI) to achieve the targeted concentration levels of NIs in the media. Recovery tests were performed in both media (SW and FW) at least at two concentration levels. Triplicate samples for each concentration level were processed. Samples were stored at laboratory temperature (25- 28°C) for at least two hours before extraction and analysis to ensure interaction of analyte and medium components.

The summarized results of the recovery tests (%) in the SW or to FW media and relative standard deviations (RSDs) (%) are shown in Table S2. The accuracy of the methods was acceptable since the average recoveries measured at the selected fortification levels were in the range of 70-120%, except for sakuranetin for which the achieved recoveries at the highest tested concentration (500 μM) were approx. 30% (28.2 and 33.7 % for SW and FW medium, respectively) due to solubility issues. The RSDs expressing the precision of the developed procedures were less than 20%, in all cases.

**Chromatographic analyses**

High performance liquid chromatography (HPLC) analyses were performed in a Shimadzu LC-20ADHPLC system equipped with an UV/VIS PDA detector. A Shimadzu GVP-ODs (4.6 mm by 150 mm, 5 mm) pre-column, connected to a RP Shimadzu VPODs (4.6 mm x 150 mm, 5 mm) column, was used for NIs separation. The injection volume was 10 μl for caffeic acid, and 20 μl for all other compounds. The flow rate of the mobile phase was set at 0.7 mL min$^{-1}$ for caffeic acid, 0.8 mL min$^{-1}$ for chlorogenic acid, simvastatin, 1,4-dibutyltriazole, and nitrapyrin, and 1 mL min$^{-1}$ for all other compounds. Column temperature was set at 40°C for 2-methoxy-1,4-naphthoquinone, and at 25 °C for all other NIs. Chromatographic separation of all compounds was achieved at isocratic conditions. Mixtures of methanol and orthophosphoric acid [0.1% (vol/vol)] were used at ratios of 40:60, 30:70 and 70:30 (vol/vol) as mobile phases for the chromatographic separation of chlorogenic acid, caffeic and shikimic acid, and sakuranetin, respectively, and detection was achieved at 325 (for chlorogenic and caffeic acid), 215 and 287 nm, respectively. Mixtures of acetonitrile and orthophosphoric acid [0.1% (vol/vol)] were used at a ratio of 80:20, 70:30, 60:40, 40:60, and 30:70 (vol/vol) as mobile phases for the chromatographic elution of simvastatin; nitrapyrin; 1-butyl-4-propyl-triazole and 1,4-dibutyltriazole; zeanone and MHPP; and 2-methoxy-1,4-naphthoquinone, respectively. Similarly, detection of these compounds was achieved at 238; 269; 220; 229; 262 and 224; and 250 nm, respectively. Table S3 summarizes the conditions used for the chromatographic separation of each compound**.** Calibration curves obtained by the injection of standard solutions of NIs in methanol or acetonitrile (in case of 2-methoxy-1,4-naphthoquinone and zeanone), ranging from 1 to 100 mg $L^{-1}$ for caffeic acid and from 0.05 to 20 mg $L^{-1}$ for all other compounds, were used for quantification. The linearity of the calibration curve was assessed by calculating the determination coefficients ($r^2$).

**Table S1.** The concentration levels (µM) used per compound for the inhibition assays.

| **Compound** | **Concentration level (µM)** | | | | | |
|---|---|---|---|---|---|---|
| Sakuranetin | 0 | 10 | 25[1] | 100 | 250 | 500[2] |
| MHPP | 0 | | 50 | 150 | 500 | 1500 |
| Zeanone | 0 | | 0.2 | 2 | 20 | 46[3] |
| 2-Methoxy-1,4-naphthoquinone | 0 | | 0.5 | 5 | 50 | 218[3] |
| Caffeic acid | 0 | 0.3[1] | 3 | 30 | 300 | 2220[2,3] |
| Quinic acid | 0 | 0.3[1] | 3 | 30 | 300 | 3000[2] |
| Chlorogenic acid | 0 | 0.15[1] | 1.5 | 15 | 150 | 1500[2] |
| Shikimic acid | 0 | | 5 | 50 | 500 | 5000 |
| Simvastatin | 0 | | 5 | 15 | 50 | 100 |
| 1-Butyl-4-propyl-triazole | 0 | | 1 | 10 | 100 | 500 |
| 1,4-Dibutyltriazole | 0 | | 1 | 10 | 100 | 500 |
| Nitrapyrin | 0 | | 5 | | | |

[1] Concentration levels used only in the bioassays with *Ca.* N. sinensis and *Ca.* N. franklandianus.
[2] Concentration levels used only in the bioassays with *N. multiformis*, *N. europaea* and *Nitrobacter* sp.
[3] Concentration levels adjusted based on the maximum solubility determined in DMSO.

**Table S2.** Recovery levels obtained for the extraction of each of the NIs tested in the different growth media used (SW: Skinner and Walker's medium; FW: MES-buffered freshwater medium). The recovery value at each concentration level is the mean of triplicates.

| Compound | Medium | Extraction method | Fortification level (µM) | Recovery (%) (Mean, n=3) | Standard deviation | Relative standard deviation | Sensitivity (%) |
|---|---|---|---|---|---|---|---|
| Sakuranetin | SW | 1:5 MeOH (vol/vol) | 500 | 28.16 | 0.31 | 1.12 | 76.49 |
| | | | 100 | 88.67 | 6.08 | 6.86 | |
| | | | 25 | 112.63 | 4.21 | 3.73 | |
| | FW | 1:5 MeOH (vol/vol) | 500 | 33.65 | 4.33 | 12.86 | 74.33 |
| | | | 100 | 86,88 | 5.00 | 5.76 | |
| | | | 25 | 102.47 | 3.01 | 2.94 | |
| MHPP | SW | 1:10 MeOH (vol/vol) | 150 | 103.95 | 2.28 | 2.19 | 107.59 |
| | | | 50 | 111.22 | 1.35 | 1.21 | |
| | FW | 1:10 MeOH (vol/vol) | 150 | 119.60 | 14.56 | 12.17 | 118.73 |
| | | | 50 | 117.85 | 15.10 | 12.81 | |
| Zeanone | SW | 1:2 MeCN (vol/vol) | 46 | 103.63 | 3.88 | 3.74 | 102.73 |
| | | | 20 | 108.05 | 4.97 | 4.60 | |
| | | | 2 | 96.50 | 5.89 | 6.11 | |
| | FW | 1:2 MeCN (vol/vol) | 46 | 74.78 | 10.64 | 14.23 | 86.79 |
| | | | 20 | 91.08 | 4.10 | 4.50 | |
| | | | 2 | 94.50 | 5.07 | 5.37 | |
| Caffeic acid | SW | 1:10 MeOH (vol/vol) | 2220 | 93.10 | 3.76 | 4.04 | 102.16 |
| | | | 300 | 99.03 | 6.40 | 6.46 | |
| | | | 30 | 114.35 | 5.76 | 5.04 | |
| | FW | 1:5 MeOH (vol/vol) | 2220 | 55.16 | 0.79 | 1.43 | 83.10 |
| | | | 300 | 98.24 | 1.36 | 1.38 | |
| | | | 30 | 95.91 | 3.50 | 3.65 | |

| | | | | | | | |
|---|---|---|---|---|---|---|---|
| **Chlorogenic acid** | **SW** | 1:5 MeOH (vol/vol) | 150 | 89.92 | 3.15 | 3.51 | 94.12 |
| | | | 15 | 90.75 | 2.02 | 2.23 | |
| | | | 1.5 | 101.69 | 3.77 | 3.70 | |
| | **FW** | 1:5 MeOH (vol/vol) | 150 | 91.64 | 2.34 | 2.56 | 100.36 |
| | | | 15 | 87.66 | 1.28 | 1.46 | |
| | | | 1.5 | 121.78 | 19.03 | 15.62 | |
| **Shikimic acid** | **SW** | 1:10 MeOH (vol/vol) | 5000 | 86.61 | 4.57 | 5.28 | 99.49 |
| | | | 500 | 90.81 | 9.96 | 10.96 | |
| | | | 50 | 121.06 | 1.65 | 1.36 | |
| | **FW** | 1:10 MeOH (vol/vol) | 5000 | 91.02 | 0.50 | 0.54 | 92.69 |
| | | | 500 | 83.90 | 2.21 | 2.63 | |
| | | | 50 | 103.14 | 0.24 | 0.23 | |
| **Simvastatin** | **SW** | 1:2 MeOH (vol/vol) | 50 | 97.53 | 3.03 | 3.10 | 95.60 |
| | | | 15 | 93.67 | 3.03 | 3.23 | |
| | **FW** | 1:10 MeOH (vol/vol) | 50 | 93.12 | 2.92 | 3.13 | 88.39 |
| | | | 15 | 83.67 | 2.54 | 3.04 | |
| **1-Butyl-4-propyl-triazole** | **SW** | 1:2 MeOH (vol/vol) | 500 | 97.42 | 1.94 | 2.00 | 97.44 |
| | | | 100 | 97.47 | 3.51 | 3.60 | |
| | **FW** | 1:2 MeOH (vol/vol) | 500 | 74.94 | 1.50 | 2.00 | 70.53 |
| | | | 100 | 66.12 | 1.91 | 2.89 | |
| **1,4-Dibutyltriazole** | **SW** | 1:2 MeOH (vol/vol) | 100 | 103.96 | 1.28 | 1.23 | 100.15 |
| | | | 10 | 96.33 | 2.89 | 3.00 | |
| | **FW** | 1:2 MeOH (vol/vol) | 100 | 100.60 | 5.94 | 5.90 | 96.67 |
| | | | 10 | 92.74 | 3.83 | 4.13 | |
| **2-Methoxy-1,4-naphthoquinone** | **SW** | 1:2 MeCN (vol/vol) | 215 | 72.26 | 0.75 | 1.04 | 81.67 |
| | | | 50 | 90.70 | 5.82 | 6.42 | |

| | | | | | | | |
|---|---|---|---|---|---|---|---|
| | | | 5 | 82.04 | 6.94 | 8.46 | |
| | **FW** | 1:2 MeCN (vol/vol) | 215 | 75.05 | 1.37 | 1.83 | 81.39 |
| | | | 50 | 90.45 | 3.07 | 3.40 | |
| | | | 5 | 78.67 | 3.50 | 4.45 | |

**Table S3.** The chromatographic conditions used for the analysis of the residues of the tested Nitrification Inhibitors (NIs).

| **Compound**[4] | **Mobile phase** | **Detection (nm)** | **Retention time (min)** | **Column Temperature ($^oC$)** | **Flow rate (ml/min)** | **Injection volume (µl)** |
|---|---|---|---|---|---|---|
| Sakuranetin | 70 MeOH : 30 $H_2O$ (0.01 %$H_3PO_4$) | 287 | 6.5 | 25 | 1.0 | 20 |
| MHPP | 40 MeCN : 60 $H_2O$ (0.01 %$H_3PO_4$) | 224 | 2.6 | 25 | 1.0 | 20 |
| Zeanone | 40 MeCN : 60 $H_2O$ (0.01 %$H_3PO_4$) | 262 | 8.4 | 25 | 1.0 | 20 |
| Caffeic acid | 30 MeOH : 70 $H_2O$ (0.01 %$H_3PO_4$) | 325 | 3.0 | 25 | 0.7 | 10 |
| Chlorogenic acid | 40 MeOH : 60 $H_2O$ (0.01 %$H_3PO_4$) | 325 | 2.6 | 25 | 0.8 | 20 |
| Shikimic acid | 30 MeOH : 70 $H_2O$ (0.01 %$H_3PO_4$) | 215 | 2.3 | 25 | 1.0 | 20 |
| Simvastatin | 80 MeCN : 20 $H_2O$ (0.01 %$H_3PO_4$) | 238 | 5.4 | 25 | 0.8 | 20 |
| 1-butyl-4-propyl-triazole | 60 MeCN : 40 $H_2O$ (0.01 %$H_3PO_4$) | 220 | 3.5 | 25 | 1.0 | 20 |
| 1,4-dibutyltriazole | 60 MeCN : 40 $H_2O$ (0.01 %$H_3PO_4$) | 220 | 5.3 | 25 | 0.8 | 20 |
| 2-methoxy-1,4-naphthoquinone | 30 MeCN : 70 $H_2O$ (0.01 %$H_3PO_4$) | 250 | 9.0 | 40 | 1.0 | 20 |
| Nitrapyrin | 70 MeCN:30 $H_2O$ (0.01 %$H_3PO_4$) | 269 | 5.0 | 25 | 0.8 | 20 |

[4]Quinic acid residues were not followed due to the inability to detect this compound in the available HPLC processing system.

### Results

#### Stability of NIs on AOM and NOB cultures

Sakuranetin in general did not follow a dose dependent degradation pattern and showed up to 60% degradation in the liquid cultures of *N. multiformis* and *N. europaea*. Similarly, in *Ca.* Nitrosotalea sinensis and *Nitrobacter* sp. liquid cultures, sakuranetin constituted at the end of the incubation period approx. 30 to 100 % of the initially recovered amount. In contrast, no significant degradation ($p > 0.05$) of sakuranetin was observed in the liquid cultures of *Ca.* N. franklandianus throughout the incubation period (Fig. S4). MHPP was significantly degraded in the liquid cultures of *N. multiformis* only at the lowest tested concentration of 50 μM, for which complete degradation was recorded. In contrast, in the liquid cultures of *N. europaea* the degradation of MHPP varied from 30 to 60% without following a dose dependent pattern. In *Ca.* Nitrosotalea sinensis liquid cultures, MHPP showed up to 40% degradation at the end of the incubation period, at the tested concentration range, while no significant degradation of MHPP was observed in the liquid cultures of *Nitrobacter sp.* Finally, in the liquid cultures of *Ca.* N. franklandianus a significant degradation of MHPP ($p < 0.05$) was observed only at the lowest tested concentrations of 50 and 150 μM for which approx. 80%, and 40% degradation was recorded, respectively (Fig. S4). Zeanone showed up to 20% degradation in the liquid cultures of *N. multiformis,* with the exception of the lowest tested concentration of 0.2 μM for which complete degradation was recorded. However, in the liquid cultures of *N. europaea* higher degradation rates of zeanone were observed, that varied from 40 to 100%. In *Ca.* Nitrosotalea sinensis liquid cultures, zeanone was significantly degraded only at the highest tested concentrations of 20 and 46 μM, constituting at 16 days approx. 40% of the initially recovered amount. In contrast in *Nitrobacter* sp. cultures zeanone was significantly degraded only at the lowest tested concentration (0.2 μM) for which complete degradation was observed. Finally, in the liquid cultures of *Ca.* N. franklandianus zeanone was completely degraded at the lowest tested concentration (0.2 μM), while the degradation for the higher tested concentrations ranged from 30- 50% (Fig. S4).

Caffeic acid and chlorogenic acid were completely degraded at all concentration levels tested in the liquid cultures of *N. multiformis* and *Ca.* N. franklandianus. Similarly, in the liquid cultures of *N. europaea* both NIs showed degradation levels > 78% for all tested concentrations. A dose dependent degradation pattern was observed for caffeic acid and chlorogenic acid in *Ca.* Nitrosotalea sinensis liquid cultures, with the NIs be fully degraded only at the lowest tested concentration levels (≤3 μM and ≤1.5 μM, respectively). Although, a similar degradation pattern was observed for caffeic acid in the liquid cultures of *Nitrobacter* sp., chlorogenic acid did not show a dose dependent degradation pattern and constituted approx. 43 to 80% of the initially recovered amount at 14 days after the onset of the incubation. Finally, shikimic acid was generally stable and significant degradation ($p < 0.05$), ranging from 85 to100%, was only observed in the liquid cultures of *N. multiformis*, *Ca.* N. franklandianus, and *Nitrobacter* sp. treated with the lowest tested concentrations (5 and 50 μM) (Fig. S5).

No significant degradation ($p > 0,05$) of the tested NI analogues was observed in the liquid cultures of *N. multiformis* throughout the incubation period, with the exception of the lowest tested concentration of 2-methoxy-1,4-naphthoquinone (0.5 μM) for which complete degradation was recorded. In the liquid cultures of *N. europaea* however, the degradation of simvastatin varied from 40 to 90% without following a dose dependent pattern. 1-butyl-4-propyl-triazole and 1,4-dibutyltriazole were generally persistent with the highest degradation rates (100% and 40% respectively) observed at the 1 μM concentration level, while for 2-methoxy-1,4-naphthoquinone approx. 70% degradation was recorded at all tested concentrations. Simvastatin was fully degraded in the liquid cultures of *Ca.* N. sinensis treated with the lower tested concentrations (15 and 5 μM), while approx. 90% degradation was recorded at the concentrations of 50 and 100 μM. On the other hand, both triazoles, and 2-methoxy-1,4-naphthoquinone were stable during the incubation period. In the liquid cultures of *Ca.* N. franklandianus simvastatin, 1-butyl-4-propyl-triazole and 1,4-dibutyltriazole showed degradation > 70% at the lowest tested concentrations of 5 μM and 1 μM, respectively. On the other hand, 2-methoxy-1,4-naphtoquinone showed 70-100% degradation at the tested concentration range. Finally, in the liquid cultures of *Nitrobacte*r sp. simvastatin showed approx. 80% degradation at the higher tested concentrations (≥ 15 μM). The degradation of 1-butyl-4-propyl-triazole did not follow a dose-dependent pattern and varied from 5 to 63%, while 1,4-dibutyltriazole showed no significant degradation throughout the incubation period. 2-methoxy-1,4-naphthoquinone, did not show significant degradation ($p > 0.05$) with the exception of its lowest tested concentration for which approx. 80% degradation was recorded (Fig. S6).

**Stability of nitrapyrin on AOM and NOB Cultures**

At the end of the incubation period nitrapyrin constituted approx. 40-50%, 1-20%, and 1% of its total amount recovered upon its addition in the liquid cultures of AOB, AOA, and NOB, respectively, at the concentration of 5 μM, in line with Papadopoulou et al. (2020) (Fig. S3).

**Table S4.** One-way analysis of strain-dependent variation in inhibitory concentration ($EC_{50}$) between AOB strains for each NI. $EC_{50}$ values of the AOB strains for each NI followed by the same letter are not significantly different at the 5% level. The asterisk(s) denotes that no $EC_{50}$ could be descent from the statistical analysis. One asterisk (*) indicates that $EC_{50}$ value is lower than the minimum tested concentration, while two asterisks (**) indicate that the $EC_{50}$ value is higher than the maximum tested concentration.

| NI | *N. multiformis* | *N. europaea* |
|---|---|---|
| Sakuranetin | 144.28 (±18.97) | >500.00** |
| MHPP | 104.03 (±12.76)b | 339.29(±32.36)a |
| Zeanone | < 0.20* | >46.00** |
| 2-Methoxy-1,4-naphthoquinone | >218.00** | >218.00** |
| Caffeic acid | 140.20 (±12.75)b | 203.29(±15.84)a |
| Quinic acid | < 3.00* | 1596.60(±123.14) |
| Chlorogenic acid | 81.15 (±9.42)b | 777.50(±49.89)a |
| Shikimic acid | 191.17 (±41.97) | >5000.00** |

| | | |
|---|---|---|
| Simvastatin | > 100.00** | > 100.00** |
| 1-Butyl-4-propyl-triazole | 407.32 (±62.98) | >500.00** |
| 1,4-Dibutyltriazole | > 500.00** | > 500.00** |

**Table S5.** One-way analysis of strain-dependent variation in inhibitory concentration ($EC_{50}$) between AOA strains for each NI. $EC_{50}$ values of the AOA strains for each NI followed by the same letter are not significantly different at the 5% level. The asterisk(s) denotes that no $EC_{50}$ could be descent from the statistical analysis. One asterisk (*) indicates that $EC_{50}$ value is lower than the minimum tested concentration, while two asterisks (**) indicate that the $EC_{50}$ value is higher than the maximum tested concentration.

| NI | *Ca.* N. sinensis | *Ca.* N. franklandianus |
|---|---|---|
| Sakuranetin | < 10.00* | < 10.00* |
| MHPP | 268.20(±3.81)b | 877.71(±165.30)a |
| Zeanone | 4.52(±0.62)a | 0.43(±0.13)b |
| 2-Methoxy-1,4-naphthoquinone | 5.22(±0.06)a | 2.92(±0.28)b |
| Caffeic acid | 1.77(±0.23)b | 19.53(±6.02)a |
| Quinic acid | 84.32(±13.98) | >300** |
| Chlorogenic acid | 65.45(±6.93)b | 148.88(±80.26)a |
| Shikimic acid | 334.87(±21.02)b | 1420.36(±167.82)a |
| Simvastatin | < 5.00* | 25.70(±5.75) |
| 1-Butyl-4-propyl-triazole | 237.37(±44.77)a | 460.03(±86.13)a |
| 1,4-Dibutyltriazole | 283.63(±71.69)a | 284.76(±46.0.3)a |

### Supplementary Figures

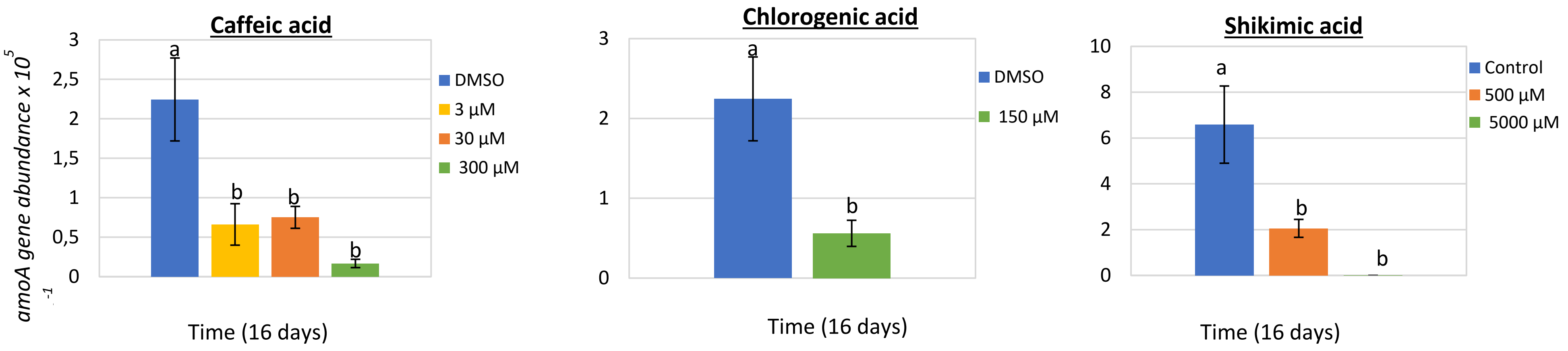


**Figure S1.** The effect of different concentrations of caffeic acid, chlorogenic acid, and shikimic acid on the growth of AOA *Ca.* N. sinensis as determined by the abundance of the *amoA* gene. Error bars represent the standard error of the mean of triplicate cultures. At each time point, bars designated with different lower-case letters are significantly different at the 5% level.

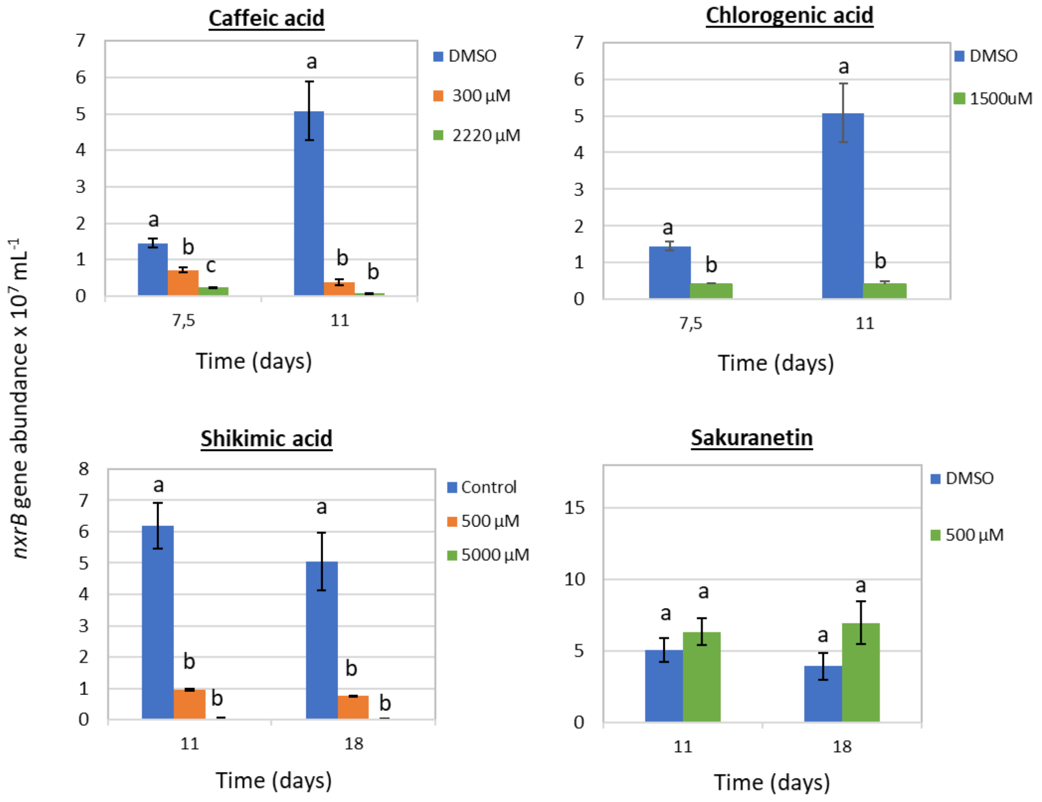


**Figure S2.** The effect of different concentrations of caffeic acid, chlorogenic acid, shikimic acid, and sakuranetin on the growth of the NOB *Nitrobacter* sp. NHB1 as determined by the abundance of the *nxrB* gene. Error bars represent the standard error of the mean of triplicate cultures. At each time point, bars designated with different lower-case letters are significantly different at the 5% level.

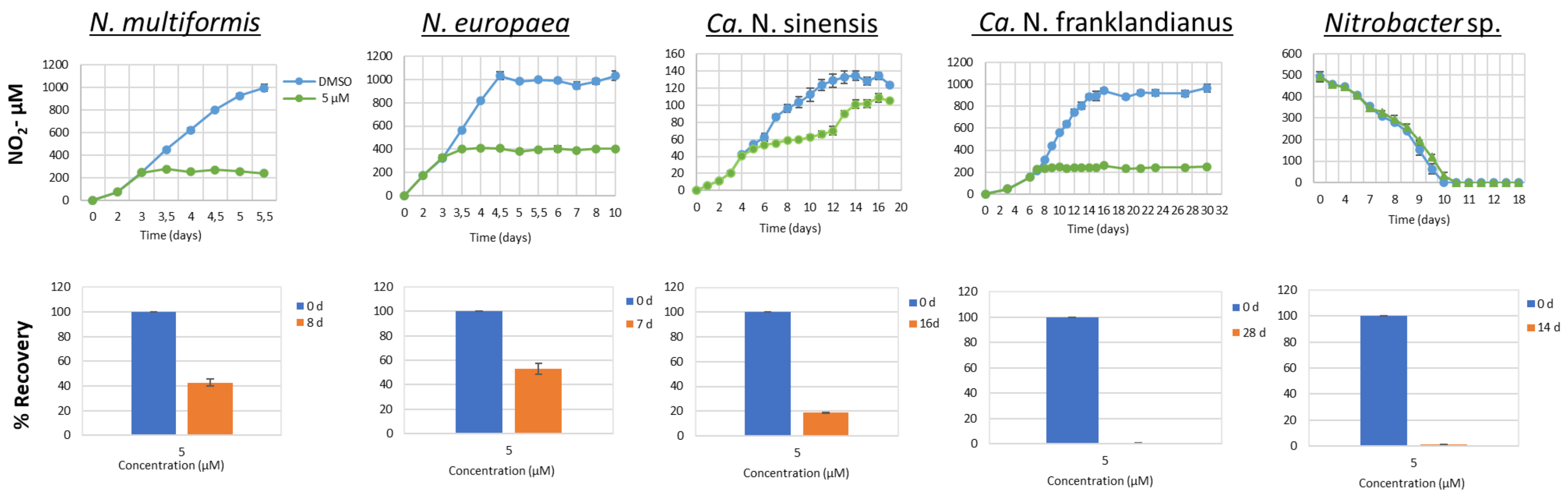


**Figure S3.** The effect of a selected concentration of nitrapyrin, used as a control of the microbial cell response, according to Papadopoulou et al. (2020), on the activity of the AOB *N. multiformis* and *N. europaea*, the AOA *Ca.* N. sinensis and *Ca.* N. franklandianus, and the NOB *Nitrobacter* sp. NHB1, as determined by monitoring the production or consumption of nitrite. The degradation of nitrapyrin is also presented. Error bars represent the standard error of the mean of triplicate cultures.

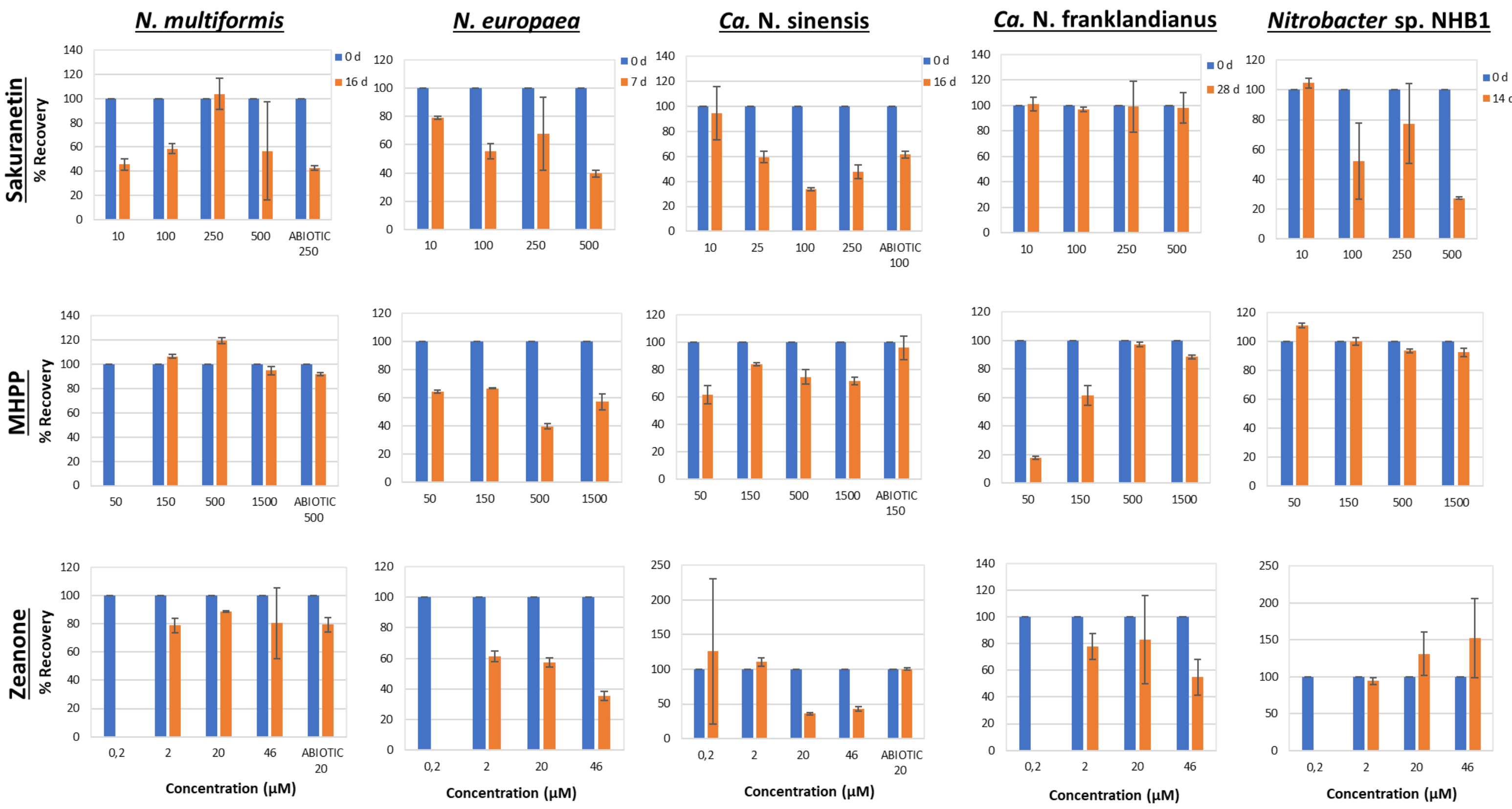


**Figure S4.** The degradation of sakuranetin, MHPP, and zeanone applied over a range of concentrations in the liquid cultures of the AOB *N. multiformis* and *N. europaea*, the AOA *Ca.* N. sinensis and *Ca.* N. franklandianus, and the NOB *Nitrobacter* sp. NHB1, and in abiotic non-inoculated controls (at one selected concentration), determined at two time points (start: early logarithmic phase, end: early stationary phase). Error bars represent the standard error of the mean of triplicate cultures.

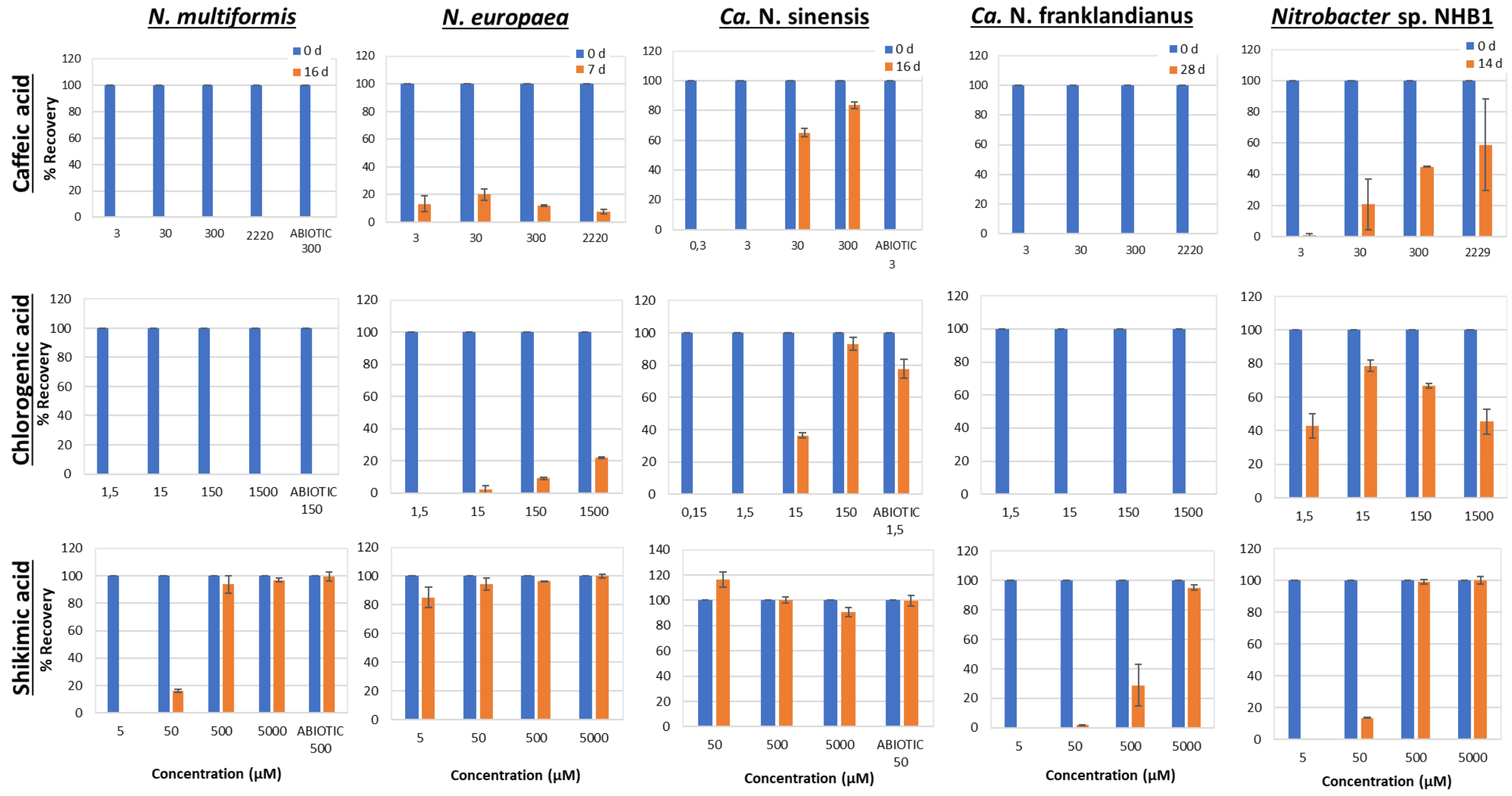


**Figure S5.** The degradation of caffeic acid, chlorogenic acid, and shikimic acid applied over a range of concentrations in the liquid cultures of the AOB *N. multiformis* and *N. europaea*, the AOA *Ca.* N. sinensis and *Ca.* N. franklandianus, and the NOB *Nitrobacter* sp. NHB1, and in non-inoculated abiotic controls (at one selected concentration), determined at two time points (start: early logarithmic phase, end: early stationary phase). Error bars represent the standard error of the mean of triplicate cultures.

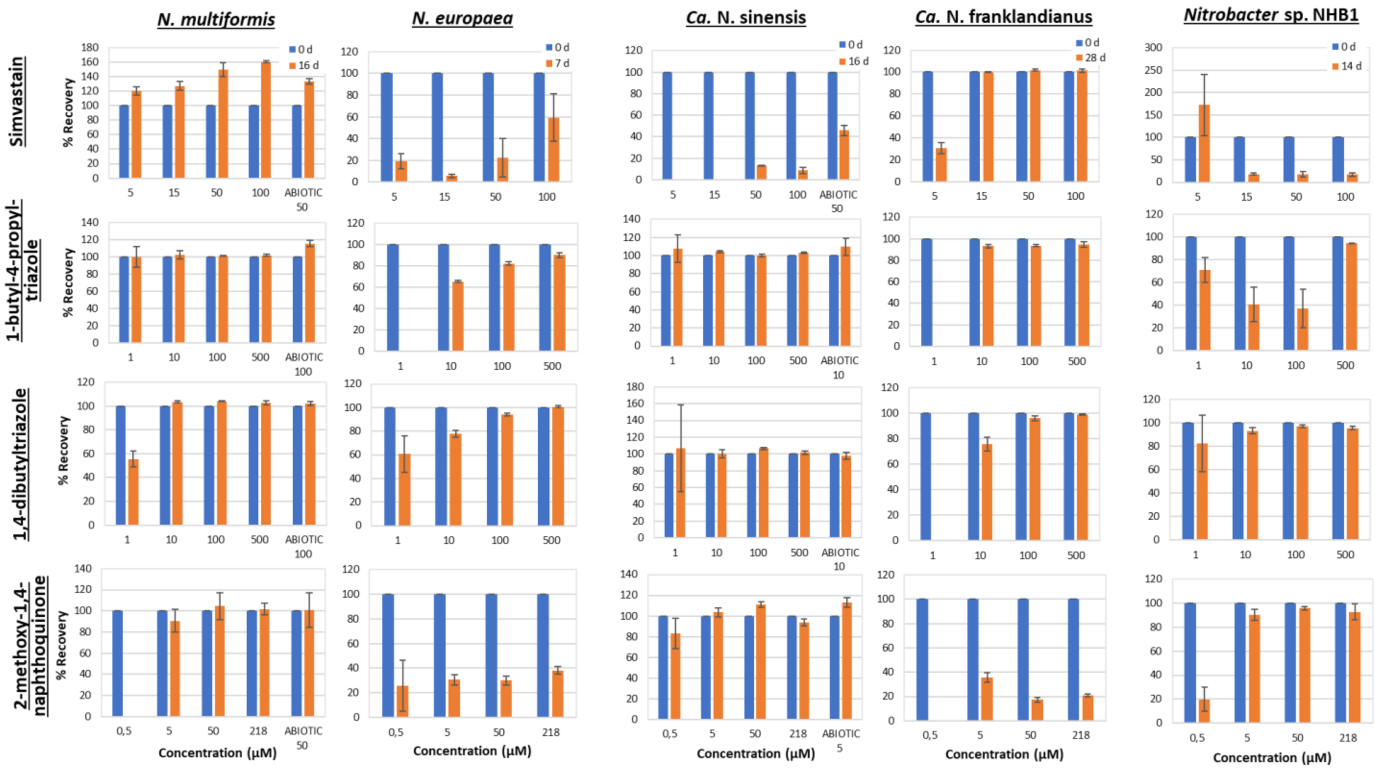


**Figure S6.** The degradation of simvastatin, 1-butyl-4-propyl-triazole, and 1,4-dibutyltriazole, and 2-methoxy-1,4-naphthoquinone applied over a range of concentrations in the liquid cultures of the AOB *N. multiformis* and *N. europaea*, the AOA *Ca.* N. sinensis and *Ca.* N. franklandianus, and the NOB *Nitrobacter* sp. NHB1, and in non-inoculated abiotic controls (at one selected concentration), determined at two time points (start: early logarithmic phase, end: early stationary phase). Error bars represent the standard error of the mean of triplicate cultures.
